# Phenotypic complexities of rare heterozygous neurexin-1 deletions

**DOI:** 10.1101/2023.10.28.564543

**Authors:** Michael B. Fernando, Yu Fan, Yanchun Zhang, Alex Tokolyi, Aleta N. Murphy, Sarah Kammourh, P.J. Michael Deans, Sadaf Ghorbani, Ryan Onatzevitch, Adriana Pero, Christopher Padilla, Sarah Williams, Erin K. Flaherty, Iya A. Prytkova, Lei Cao, David A. Knowles, Gang Fang, Paul A. Slesinger, Kristen J. Brennand

## Abstract

Given the large number of genes significantly associated with risk for neuropsychiatric disorders, a critical unanswered question is the extent to which diverse mutations --sometimes impacting the same gene--will require tailored therapeutic strategies. Here we consider this in the context of rare neuropsychiatric disorder-associated copy number variants (2p16.3) resulting in heterozygous deletions in *NRXN1*, a pre-synaptic cell adhesion protein that serves as a critical synaptic organizer in the brain. Complex patterns of *NRXN1* alternative splicing are fundamental to establishing diverse neurocircuitry, vary between the cell types of the brain, and are differentially impacted by unique (non-recurrent) deletions. We contrast the cell-type-specific impact of patient-specific mutations in *NRXN1* using human induced pluripotent stem cells, finding that perturbations in *NRXN1* splicing result in divergent cell-type-specific synaptic outcomes. Via distinct loss-of-function (LOF) and gain-of-function (GOF) mechanisms, *NRXN1^+/-^* deletions cause decreased synaptic activity in glutamatergic neurons, yet increased synaptic activity in GABAergic neurons. Reciprocal isogenic manipulations causally demonstrate that aberrant splicing drives these changes in synaptic activity. For *NRXN1* deletions, and perhaps more broadly, precision medicine will require stratifying patients based on whether their gene mutations act through LOF or GOF mechanisms, in order to achieve individualized restoration of *NRXN1* isoform repertoires by increasing wildtype, or ablating mutant isoforms. Given the increasing number of mutations predicted to engender both LOF and GOF mechanisms in brain disorders, our findings add nuance to future considerations of precision medicine.

## MAIN

Neurexins are pre-synaptic cell adhesion proteins that act as synaptic organizers^1^. There are three neurexin genes (*NRXN1*, *NRXN2*, and *NRXN3*) in mammals and each is highly alternatively spliced to produce hundreds of isoforms, primarily categorized into long alpha and shorter beta isoforms^2^. The complex alternative splicing of neurexins expands protein-protein interaction capabilities^3^, allowing neurexins to interact with diverse post-synaptic ligands to establish and maintain neurotransmission. *NRXN1α* splice variants are specific to brain regions^4^ and between cell types^5^, but the cell-type-specific functional impact of individual isoforms remains unclear. Although rare in the population, large copy number variations (deletions or duplications) at the *NRXN1* locus 2p16.3, particularly those deleting exonic regions, are highly penetrant, pleiotropic, and are strongly associated with several neuropsychiatric diseases, including schizophrenia (odds ratio 14.4^6^), autism spectrum disorder (odds ratio 14.9^7^), epilepsy (odds ratio 9.91^8^), intellectual disability (odds ratio 7.47^9^) and Tourette’s syndrome (odds ratio 20.3^10^). Deletions in *NRXN1* are non-recurrent (that is, they vary in size and location), making it difficult to determine the molecular mechanisms underlying their diverse clinical symptoms (e.g., diagnosis, severity, prognosis, and age-of-onset). In rodent studies, gene knockouts (KO) of *NRXNs* are sufficient to produce an array of excitatory and inhibitory synaptic phenotypes^1^. However, heterozygous deletions yield only modest behavioral and physiological changes *in vivo*^11^. By contrast, *in vitro* studies of engineered heterozygous *NRXN1*^+/-^ human neurons reveal robust changes in excitatory neurotransmission that are not recapitulated in matched *NRXN1*^+/-^ mouse neurons^12,13^, and studies that utilize patient derived *NRXN1*^+/-^ neurons have yet to deconvolute complex phenotypes in a cell-type and genotype specific manner^12,14^. Altogether, neurexins possess unique human neurobiology, and therefore, the impact of distinct patient-specific *NRXN1^+/-^* must be specifically evaluated in human models with an important consideration of how unique *NRXN1^+/-^* deletions impact splicing patterns and neuronal function across distinct cell-types.

Human induced pluripotent stem cell (hiPSC)-derived neurons provide an ideal platform to study *NRXN1α* alternative splicing. Previously we established that hiPSC-derived forebrain cultures, comprised of a mixture of glutamatergic and GABAergic neurons with astroglia, recapitulate the diversity of *NRXN1α* alternative splicing observed in the human brain, cataloguing 123 high-confidence *NRXN1α* isoforms^15^. Furthermore, using patient-derived *NRXN1^+/-^* hiPSCs with unique 5’- or 3’-deletions in the gene, we uncovered wide-scale reduction in wildtype *NRXN1α* isoform levels and, robust expression of dozens of novel isoforms from the 3’-deletion allele only^15^. Overexpression of individual wildtype isoforms (WT) ameliorated reduced neuronal activity in patient-derived *NRXN1^+/-^* hiPSC-neurons in a genotype-dependent manner, whereas mutant isoform (MT) expression decreased neuronal activity levels in control hiPSC-neurons^15^. We therefore hypothesized that 5’-deletions of the promoter region represent classical loss-of-function (LOF), while robust expression of novel 3’-specific MT isoforms confer a gain-of-function (GOF) effect that cannot be rescued by overexpression of WT isoforms. Although *NRXN1* splicing varies between the cell types of the brain, the impact of non-recurrent *NRXN1^+/-^* deletions on cell-type-specific splicing patterns and synaptic function remains untested.

Neurexin signaling impacts both glutamatergic and GABAergic synapse properties^3,16^, suggesting that neurexins may regulate excitatory and inhibitory balance, which is strongly implicated across neuropsychiatric disorders^17^. Given its multifaceted roles, the lack of mechanistic understanding of how aberrant *NRXN1* splicing impacts neuronal physiology in a cell-type *and* genotype-dependent manner presents a significant challenge for therapeutic targeting. To discover the disease mechanisms that underpin *NRXN1^+/-^*deletions, we compared excitatory and inhibitory neurons, across LOF and GOF deletions, to specifically evaluate cell-autonomous phenotypes arising from distinct *NRXN1^+/-^* deletions. We identified points of phenotypic divergence across glutamatergic and GABAergic neurons, which were independently validated in isogenic experiments, thereby establishing causal relationships between aberrant splicing and synaptic dysfunction. Finally, we evaluated novel therapeutic agents for *NRXN1^+/-^*deletions based on LOF/GOF stratified patient mechanisms.

## RESULTS

### Patient-specific *NRXN1^+/-^* mutations produces differential *NRXN1* splicing patterns across glutamatergic and GABAergic neurons

To examine the impact of *NRXN1^+/-^* deletions in two different neuronal cell types, we used transcription-factor based lineage conversion of hiPSCs to generate excitatory glutamatergic neurons or inhibitory GABAergic neurons from our previous cohort^14^ (all available clinical and experimental information reported in **Supplementary Table 1**). Two cases sharing a ∼115-kb deletion in the 5’-region of *NRXN1* (5’-Del), affecting the alpha-transcript promoter, represent the LOF condition. Two additional cases sharing a ∼136-kb deletion in the 3’-region of *NRXN1* (3’-Del), impacting two alternative splice sites (SS4, SS5) across three exons (21-23), robustly express unique MT isoforms from the affected allele and therefore represent the GOF condition^15^. For controls, we used passage-matched hiPSCs from four healthy sex-balanced subjects (**Fig. 1a,b**). Transient overexpression of *NGN2* in hiPSCs produced iGLUT neurons that were >95% pure glutamatergic neurons, robustly expressed glutamatergic genes, released glutamate, and produced spontaneous synaptic activity by day 21 in vitro^18,19^. On the other hand, transient overexpression of *ASCL1* and *DLX2* yielded iGABA neurons that were >80% positive for expressing GABA and GAD1/2 by day 35 in vitro, and possessed mature physiologic properties of inhibitory neurons by day 42^20,21^. Immunostaining confirmed expression of neurotransmitter transporters, vGLUT1 and vGAT for iGLUT and iGABA neurons, respectively (**Fig. 1c,h**). RNA-sequencing (RNAseq) further validated iGLUT (DIV21) and iGABA (DIV35) neuronal induction in all donors (**Fig. 1d,i**).

**Figure 1.**
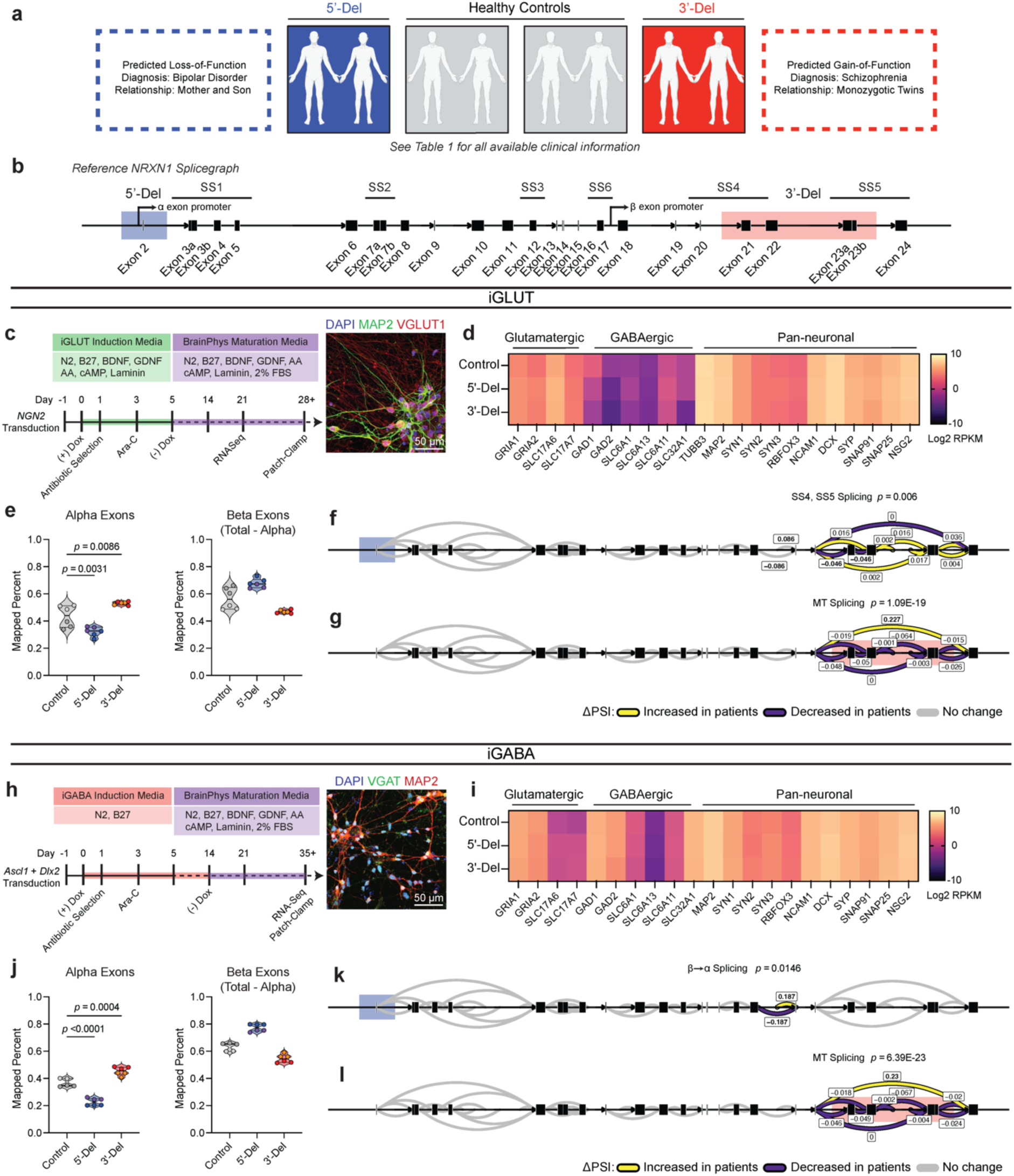
*Aberrant NRXN1 splicing across human iPSC derived glutamatergic (iGLUT) and GABAergic (iGABA) neurons.* (**a**) Brief description of clinical information of all hiPSC lines used in this study, and (**b**) schematic of *NRXN1* gene structure as a splicegraph, denoting splice sites (SS1-6), with red and blue shades corresponding on 3’-Del and 5’-Del genotypes, respectively. Arrows indicate relative internal promoter positions. (**c, h**) Induction timeline and factors to generate iGLUT and iGABA neurons, with immunostaining validation of neuronal identity (MAP2), glutamate identity (vGLUT1), and GABA identity (vGAT). (**d, i**). Gene expression panel confirming abundance of neuronal markers and neurotransmitter identity in iGLUT (n: Control = 6/2; 5’-Del = 6/2; 3’-Del = 6/2 | 1 batch) and in iGABA neurons (n: Control = 5/2; 5’-Del = 6/2; 3’-Del = 6/2 | 1 batch). (**e, j**) Mapped percent of alpha *NRXN1* exon reads in iGLUT, compared via a 1-way ANOVA, with Dunnett’s test (F_2, 15_ = 25.70; 5’-Del p = 0.0031, 3’-Del p = 0.0086), and iGABA neurons (F_2, 14_ = 92.92; 5’-Del p < 0.001, 3’-Del p = 0.0004), with beta *NRXN1* exon reads, calculated by subtracting alpha-specific reads against total reads. (**f, k**) Splicegraphs displaying significant gene wide splicing clusters, compared via Dirichlet-multinomial generalized linear model with Bonferroni corrections, for 5’-Del iGLUT (SS4 and SS5 cluster *p* = 0.006) and iGABA neurons (β→α cluster *p* = 0.0146), and for (**g, l**) 3’-Del iGLUT (SS4 and SS5 cluster *p* = 1.09E-19) and iGABA neurons (SS4 and SS5 cluster *p* = 6.39E-23). n reported as samples/donors | independent batches.

Despite cell-type-specific *NRXN1* isoform profiles reported in healthy brains^5^ and neurons^15^, aberrant *NRXN1* alpha/beta exon expression patterns appeared similar between *NRXN1^+/-^* iGLUT and iGABA neurons (**Fig. 1e,j**), whereby alpha exon expression (exons 2-18) decreased for 5’-Del and increased for 3’-Del in both types of neurons (iGLUT 5’-Del *p* = 0.0031; 3’-Del, *p* = 0.0086, 1-way ANOVA, Dunnett’s test; iGABA 5’-Del *p* = 0.0004; 3’-Del, *p* < 0.0001, 1-way ANOVA, Dunnett’s test). To investigate changes in alternative splicing, we applied LeafCutter^22^, an annotation free analysis to estimate differential splicing events visualized by splicegraphs denoting change of percent-spliced-in (ΔPSI) ratios across the *NRXN1* gene at annotated splice sites (SS1-6) and novel unannotated junctions (**Fig 1b**). Differential splicing analysis between iGLUT and iGABA neurons produced a single significant cluster (out of nine identified), centered around SS3 (**Extended Data Fig. 1a**, p = 0.01960, Bonferroni corrected), suggesting that the majority of isoforms were conserved between the two cell-types, as previously reported^15^.

Specifically, we detected both canonical and noncanonical splicing patterns in 5’-Del and 3’-Del neurons, relative to controls. Splicing of *NRXN1* transcript was perturbed at two locations in 5’-Del neurons. The first change in splicing occurred at the alternative transcription start site, reducing splicing of beta-exon transcripts into alpha-exon transcripts (henceforth referred to as β→α), as expected given the affected alpha promoter. We observed a significant reduction in 5’-Del iGABA neurons (ΔPSI −0.187, *p =* 0.0146, Bonferroni corrected), and a more modest decrease in iGLUT neurons (ΔPSI −0.086, *p =* 0.1821, Bonferroni corrected), consistent with a LOF. In 5’-Del iGLUT neurons, we also detected differential splicing at SS4 inclusion (ΔPSI −0.046, *p =* 0.006, Bonferroni corrected) (**Fig. 1f,k**). For 3’-Del neurons, we observed a robust increase in a mutant splice junction between exon 20-24 in iGLUT (ΔPSI 0.227, *p* = 1.09E-19) and iGABA (ΔPSI 0.23, *p* = 6.93E-23, Bonferroni corrected) neurons, concurrent with reduced wildtype splicing around SS4 and SS5 (**Fig. 1g,l**), consistent with a GOF. Interestingly, STAR-family RNA-binding proteins that regulate *NRXN1* splicing at SS4^23–26^ were also dysregulated across cell-types and genotypes **Extended Data Fig. 1b,f**). Taken together, these differential splicing patterns support our hypothesis that 5’-Del and 3’-Del mutations confer LOF/GOF phenotypes, either by reducing β→α splicing or producing novel MT isoforms, respectively.

### *NRXN1^+/-^* mutations broadly impact synaptic and neurodevelopmental pathways in induced and organoid-derived glutamatergic and GABAergic neurons

To unbiasedly evaluate the transcriptomic impact of *NRXN1^+/-^* deletions, and their unique splicing patterns, we utilized differential expression analysis^27^ in iGLUT and iGABA neurons, which revealed overlap of FDR-corrected differentially expressed genes (DEGs) across cell types/genotypes (e.g., *FAM66D, TTC34, GALNT9*), and represented gene sets related to synaptic function (via SynapseGO^28^) (**Fig. 2a-d, j-m**). To test for pathway enrichment among top DEGs (filtered by ±1.5 Log_2_FC), we performed gene-ontology analyses via ClusterProfiler^29^, against a background of all expressed genes to avoid exclusion of biologically relevant low abundance genes. 5’-Del neurons revealed a robust enrichment of terms related to neurotransmission and synaptic function in iGLUT neurons, and ligand-gated anion channel activity in iGABA neurons (**Supplementary Table 2**). 3’-Del neurons were enriched for terms related to neurodevelopment, significant in iGABA neurons, but only nominally enriched in iGLUT neurons. DEGs were distinct between genotypes (**Extended Data Fig. 1c,d,g,h**), with different DEGs in hierarchical clustered clades enriched for DNA-binding related GO terms (**Extended Data Fig. 1e,i).** Risk enrichments for schizophrenia, bipolar disorder, and autism spectrum disorder^30,31^ were most enriched in 5’-Del iGLUT neurons and 3’-Del iGABA neurons (**Extended Data Fig. 2**).

**Figure 2:**
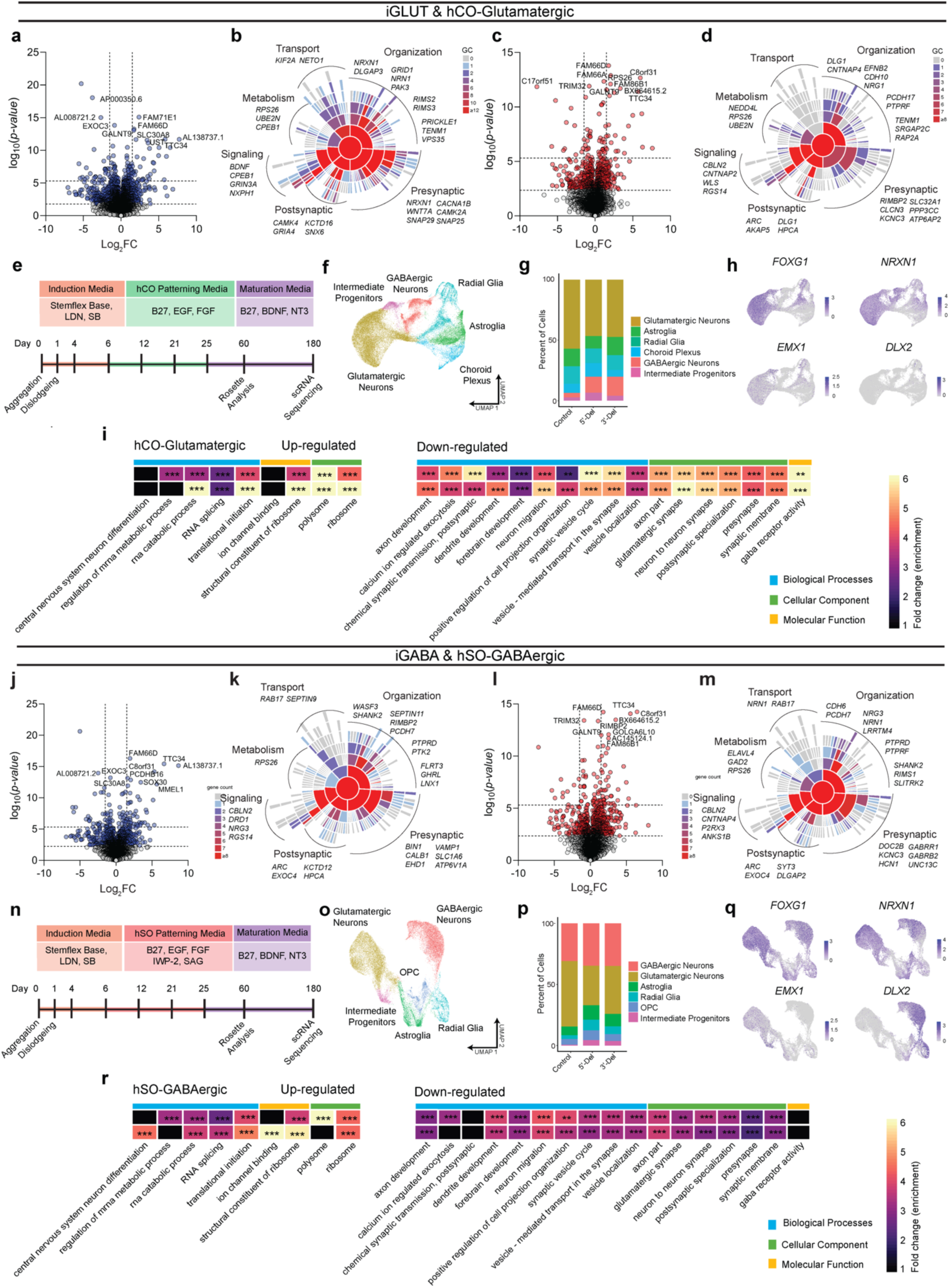
*Transcriptomic impact of NRXN1^+/-^ in induced (iGLUT/iGABA) and organoid-derived hCO-glutamatergic and hSO-GABAergic neurons.* (**a, c**) Volcano plots of differential gene expression (DE) analysis across both genotypes in iGLUT and (**j, l**) iGABA neurons. Vertical dashed lines represent DE genes ±1.5 Log2FC. Horizontal dashed lines represent FDR = 0.1 cutoff (lower) and Bonferroni corrected cutoff (upper). (**b, d**) Sunburst plots of all FDR corrected DEGs with SynGO annotated synapse function for iGLUT and (**k, m**) iGABA neurons. (**e, n**) Timeline of neural organogenesis for hCO and hSOs (**f, o**), UMAPs of hCO and hSO organoid samples sequenced at 6 months, annotated by cell clusters, and (**g, p**) relative proportions of cell clusters across genotypes, (hCO = 47,460 cells) and (hSO = 35,563 cells). (**h, q**) validation of regionalization across forebrain (*FOXG1*), dorsal (*EMX1*) and ventral (*DLX2*) regions, with *NRXN1* expression across all cells. (**i, r**) Gene ontological analysis results using DEGs from scRNASeq. *\*P* < 0.05, *\*\*P* < 0.01, *\*\*\*P* <0.001, Wilcoxon’s rank sum test, FDR = 0.05. Data represented as mean ± sem. n reported as samples/donors | independent batches.

To explore aberrant *NRXN1* splicing in a more complex neurodevelopmental system, we studied the effect of 5’-Del and 3’-Del in organoids. We applied dorsal forebrain patterning to yield human cortical organoids (hCOs) that resembled the pallium, or applied ventral forebrain patterning to generate human subpallial organoids (hSOs) that resembled the subpallium (**Fig. 2e,n and Extended Data Fig. 3a-c,h-j**)^32–34^. To stratify LOF and GOF genotypes, we confirmed exclusive MT *NRXN1* isoforms in 3’-Del organoids (**Extended Data Fig. 3d-e, k-l**). We performed single cell RNA-seq (n = 47,460 cells from hCOs, n= 35,563 cells from hSOs using 10x genomics) at 6 months, a timepoint with well characterized neural activity^33,35^, and subsequently identified clusters of cell-types within hCOs and hSOs (**Fig.2f,o**), without significant differences in cell frequencies across pooled genotypes (quasibinomial regression model **Fig. 2g,p**). Differentially expressed gene sets in hCO-glutamatergic and hSO-GABAergic clusters identified robust enrichment of GO terms related to RNA splicing (from upregulated genes), and neurodevelopment and synaptic function (from downregulated genes) (**Fig. 2h,i,q,r**). Taken together, these results reinforce the hypothesis that perturbations in *NRXN1* splicing converge on synaptic function^14,36^.

### Patient-specific alterations in spontaneous neural activity occur with minimal changes in passive and excitable membrane properties

To evaluate the functional consequence of *NRXN1^+/-^* deletions in iGLUT and iGABA neurons, we next conducted a population-level analysis of spontaneous neuronal activity using a multi-electrode array (MEA) and an examination of passive membrane properties using whole-cell patch-clamp electrophysiology. Spontaneous network activity (weighted mean firing rate, wMFR) in both 5’- and 3’-Del *NRXN1^+/-^* iGLUT neurons, which increased over time in a linear fashion (**Fig 3a**), was reduced by over 40% across two independent time points, at weeks post induction (WPI)4 (**Fig. 3b** 5’-Del *p* = 0.0005; 3’-Del, *p* < 0.0001, 1-way ANOVA, Dunnett’s test) and WPI6 (**Fig 3c** 5’-Del *p* = 0.0129, 3’-Del *p =* 0.0069 1-way ANOVA, Dunnett’s test). The cell capacitance, membrane resistance and resting membrane potentials did not significantly differ between 5’- and 3’-Del cases and control iGLUT neurons (**Fig. 3d**). To examine the intrinsic excitability, we compared the input-output curve for induced firing and found slightly lower firing in 5’-Del iGLUT neurons, but no difference in 3’-Del iGLUT neurons (**Fig. 3e**). The voltage-dependent sodium and potassium current densities were similar across 5’-Del, 3’-Del and control iGLUT neurons (**Extended Data Fig. 4a**). Taken together, these results suggest that changes in passive or intrinsic excitability membrane properties cannot fully explain the reduced firing observed on MEAs for 5’-Del and 3’-Del in iGLUT neurons.

**Figure 3.**
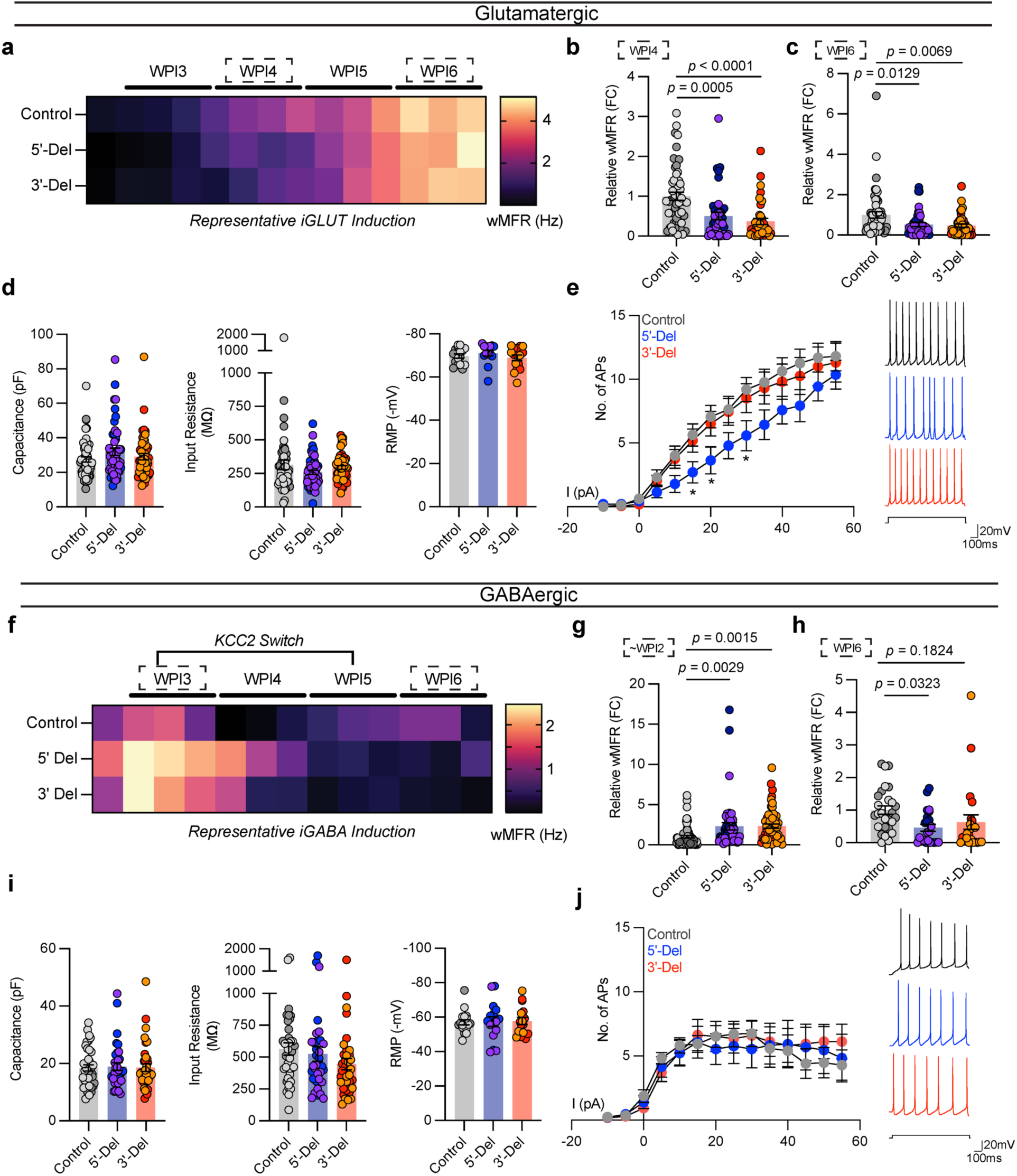
*Spontaneous, passive, and excitable properties are minimally changed from NRXN1^+/-^ induced neurons.* (**a, f**) Timelapse of multi-electrode array recordings every 2-3 days apart starting near ∼DIV12 for a single representative induction. Tiles represent averaged wMFR values across genotypes during a single recording session. (**b**) MEA quantification of iGLUT neuronal activity, compared via a 1-way ANOVA, with Dunnett’s test (n: Control = 52/2; 5’-Del = 42/2; 3’-Del = 38/2 | 3 batches) at WPI4 (F_2, 129_ = 12.77; 5’-Del *p* = 0.0005, 3’-Del *p <* 0.0001), and (**c**) WPI6 (F_2, 129_ = 5.737; 5’-Del *p* = 0.0129, 3’-Del *p =* 0.0069). (**d**) intrinsic properties of iGLUT neurons (n: Control = 51/2; 5’-Del = 51/2; 3’-Del = 48/2 | 8 batches): compared via 1-way ANOVA, with Dunnett’s test, Capacitance (F_2, 147_ = 1.505; 5’-Del *p =* 0.1565, 3’-Del *p =* 0.7633), Input resistance (F_2, 147_ = 1.311; 5’-Del *p =* 0.1949, 3’-Del *p =* 0.7906), and RMP (n: Control = 16/2; 5’-Del = 14/2; 3’-Del = 16/2 | 2 batches) compared via 1-way ANOVA, with Dunnett’s test (F_2, 43_ = 0.8234; 5’-Del *p =* 0.6378, 3’-Del *p =* 0.8468). (**e**) Input-output curves of excitable properties (n: Control = 16/2; 5’-Del = 14/2; 3’-Del = 16/2 | 2 batches), with representative traces (right), compared via Step x Genotype 2-way ANOVA; Dunnett’s Test (F_26, 559_ = 1.690, 5’-Del *p <* 0.01 at Step 6, 7 and 9, 3’-Del *p =* n.s. on all steps). (**g**) MEA quantification of iGABA neuronal activity, compared via a 1-way ANOVA, with Dunnett’s test (n: Control = 71/2; 5’-Del = 48/2; 3’-Del = 55/2 | 5 batches) at WPI2 (F_2, 171_ = 7.805; 5’-Del *p* = 0.0029, 3’-Del *p =* 0.0015), and (**h**) WPI5 (n: Control = 27/4; 5’-Del = 23/2; 3’-Del = 23/2 | 2 batches) compared via 1-way ANOVA, with Dunnett’s test (F_2, 70_ = 3.158; 5’-Del *p* = 0.0323, 3’-Del *p =* 0.1824). (**i**) intrinsic properties of iGABA neurons (n: Control = 39/2; 5’-Del = 34/2; 3’-Del = 37/2 | 6 batches): compared via 1-way ANOVA, with Dunnett’s test, Capacitance (F_2, 107_ = 0.04256; 5’-Del *p =* 0.9425, 3’-Del *p =* 0.9941), Input resistance (n: Control = 39/2; 5’-Del = 34/2; 3’-Del = 36/2 | 6 batches, F_2, 106_ = 1.451; 5’-Del *p =* 0.8162, 3’-Del *p =* 0.1703) and RMP (n: Control = 18/2; 5’-Del = 16/2; 3’-Del = 16/2 | 2 batches) compared via 1-way ANOVA, with Dunnett’s test (F_2, 47_ = 0.02842; 5’-Del *p =* 0.9893, 3’-Del *p =* 0.9601). (**j**) Input-output curves of excitable properties (n: Control = 16/2; 5’-Del = 14/2; 3’-Del = 16/2 | 2 batches), with representative traces (right), compared via Step x Genotype 2-way ANOVA; Dunnett’s Test (F_26, 572_ = 0.5137, 5’-Del and 3’-Del *p =* n.s. on all steps). Data represented as mean ± sem. n reported as samples/donors | independent batches.

In parallel, we generated iGABA neurons with the same 5’-Del, 3’-Del, and control hiPSCs, as above. We observed that immature *NRXN1^+/-^* iGABA neurons exhibited a robust ∼2-fold increase in population-wide wMFR activity (∼WPI2) from both 5’-Del and 3’-Del cases (5’-Del *p* = 0.0029, 3’-Del *p =* 0.0015 1-way ANOVA, Dunnett’s test) (**Fig. 3f,g**). Though unexpected for GABA neurons, the finding is consistent with activation of ionotropic GABA receptors leading to depolarization due to low KCC2 expression and high chloride levels^37^. Indeed, immature iGABA neurons expressed higher (10-fold increase compared to WPI2 neurons) levels of *SLC12A5* (the gene encoding *KCC2*) (**Extended Data Fig. 5a** *p* < 0.0001, 2-way ANOVA, Dunnett’s test). Furthermore, this transient hyperexcitability was pharmacologically inhibited by 10μM gabazine, a selective GABA_A_ antagonist (**Extended Data Fig. 5b,c**). In mature iGABA neurons (WPI6), however, the average wMFR decreased in 5’-Del and 3’-Del neurons (5’-Del *p* = 0.0323; 1-way ANOVA, Dunnett’s test) (**Fig. 3h**). Like iGLUT neurons, the passive and intrinsic excitability membrane properties of mature iGABA neurons were not different between *NRXN1^+/-^* 5’-Del or 3’-Del and controls (**Fig. 3i,j and Extended Data Fig. 4d**). Overall, these data suggest that patient-specific changes in spontaneous neural activity are not fully explained by differences in passive and excitable membrane properties in iGLUT or iGABA neurons from *NRXN1^+/-^* 5’-Del and 3’-Del patients.

### *NRXN1*^+/-^ 5’ and 3’ deletions result in divergent synaptic transmission deficits

To further dissect the factors mediating phenotypes in altered spontaneous firing, we investigated the efficacy of synaptic transmission by patch-clamp electrophysiology. Voltage-clamp recordings of spontaneous excitatory post-synaptic currents (sEPSCs, no TTX) in iGLUT neurons revealed decreased frequency of events for both for 5’-Del and 3’-Del iGLUT neurons (**Fig. 4a**). The cumulative probabilities of inter-event-intervals (IEI) for both 3’-Del (*p* = 4.88E-5, Levene’s test with Bonferroni correction) and 5’-Del iGLUTs (*p* = 2.54E-11, Levene’s test with Bonferroni correction) (**Fig. 4a,b**) was significantly increased, compared to controls. The sEPSC amplitude increased for 5’-Del neurons (*p* = 3.88E-4, Levene’s test with Bonferroni correction) (**Fig. 4a,c**). Miniature excitatory post-synaptic currents (mEPSCs, +TTX) showed similar trends in IEI but no changes in amplitude sizes across genotypes (IEI 5’-Del *p* = 0.0411, 1-way ANOVA, Dunnett’s test) (**Extended Data Fig. 4b,c**). These reductions in synaptic transmission are consistent with the transcriptomic signatures; pre-synaptic (SynGO) genes showed a larger change (than post-synaptic) in synaptic gene expression signatures (5’-Del Pre-SynGO, Log_2_FC = 0.1252; 3’-Del Pre-SynGO, Log_2_FC= −0.0156) (**Fig. 4d,e**). We further probed transcriptional signatures of known *NRXN1* trans-synaptic interaction partners that mediate synapse formation, function, plasticity and are frequently linked with neuropsychiatric disease^16^, and found they were represented in DEGs, including *CBLN2*, *LRRTM4* and *NXPH1*, suggesting these synaptic effects are largely driven by change in *NRXN1* expression (**Fig. 4f**).

**Figure 4:**
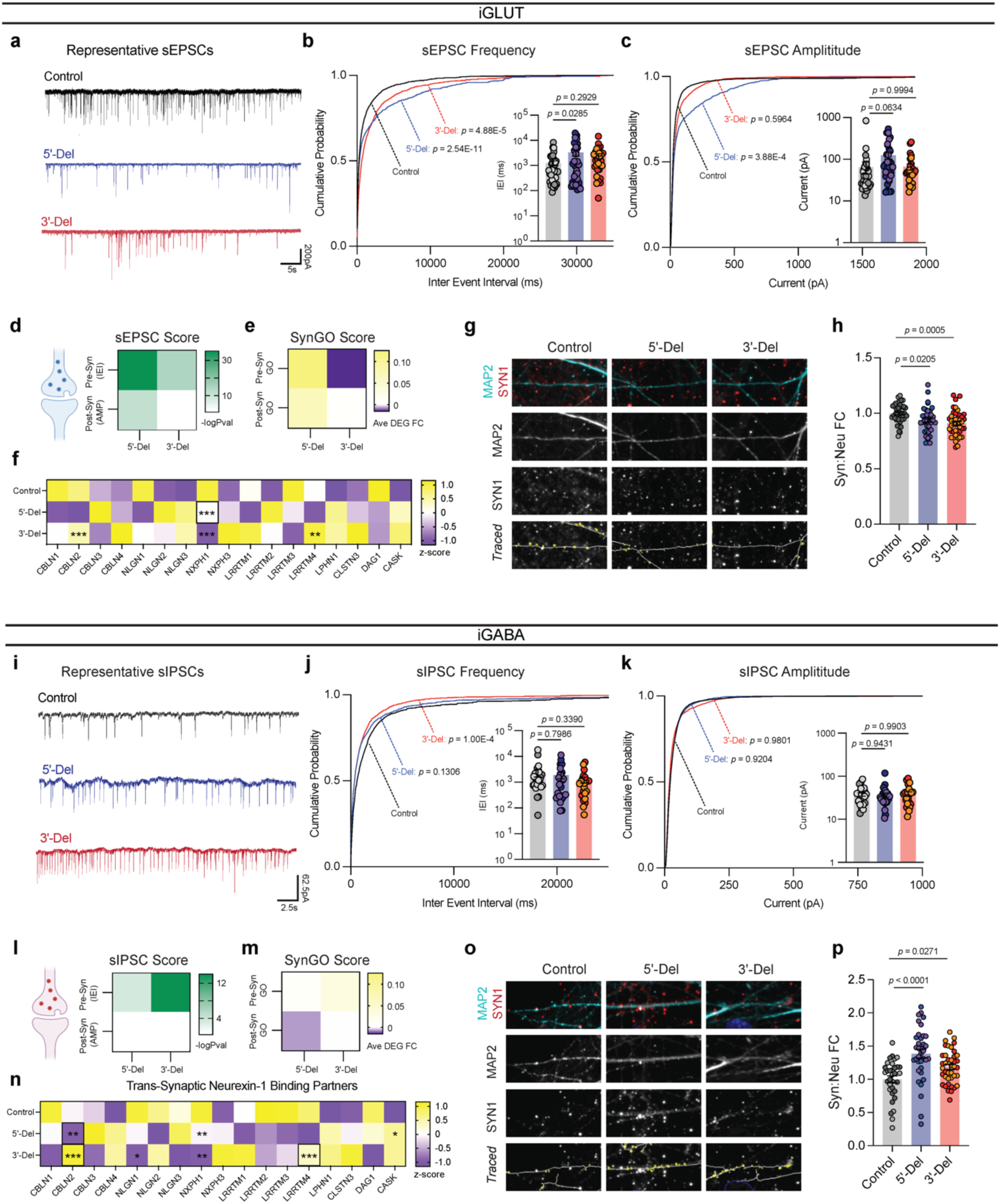
*Divergent impact on neurotransmission from NRXN1^+/-^ induced neurons.* (**a**) Representative traces of iGLUT sEPSCs. (**b**) Cumulative probabilities and log-scaled cell averages of inter-event-internals (IEIs) across genotypes (n: Control = 39/4; 5’-Del = 33/3; 3’-Del = 29/2 | 6), compared by Levene’s Test with Bonferroni correction for averaged distributions (5’-Del F = 46.635, df = 1, *p* = 2.54E-11; 3’-Del F = 17.928, df = 1, *p* = 4.88E-5), and 1-way ANOVA, with Dunnett’s test for inset (F_2, 98_ = 3.117; 5’-Del *p =* 0.0285, 3’-Del *p =* 0.2929). (**c**) Cumulative probabilities and log-scaled cell averages of amplitude size across genotypes, compared by Levene’s Test with Bonferroni correction for averaged distributions (5’-Del F = 14.15, df = 1, *p* = 3.88E-4; 3’-Del F = 1.0851, df = 1, *p* = 0.5964), and 1-way ANOVA, with Dunnett’s test for inset (F_2, 98_ = 2.839; 5’-Del *p* = 0.0634, 3’-Del *p =* 0.9994). (**d**) Transformed p-values of Levene’s Test and (**e**) SynGO gene-set averaged log_2_FC values across pre- or post-synaptic genes. (**f**) Gene expression panel (z-scores), of canonical *NRXN1* binding partners in iGLUT neurons, boxes indicate reaching genome wide significance. (**g**) Representative images of iGLUT synaptic puncta traced (SYN1) onto dendrites (MAP2), and (**h**) normalized fold change of SYN1 puncta to length of MAP2 ratio (Syn:Neu), (n: Control = 40/2; 5’-Del: 39/2; 3’-Del: 40/2), compared via 1-way ANOVA, with Dunnett’s test (F_2, 116_ = 7.538, 5’-Del *p =* 0.0205, 3’-Del *p =* 0.0005). (**i**) Representative traces of iGABA sIPSCs. (**j**) Cumulative probabilities and log-scaled cell averages of inter-event-internals (IEIs) across genotypes (n: Control = 26/2; 5’-Del = 25/3; 3’-Del = 22/2 | 4), compared by Levene’s Test with Bonferroni correction for averaged distributions (5’-Del F = 3.4002, df = 1, *p* = 0.1306; 3’-Del F = 16.501, df = 1, *p* = 1.00E-04), and 1-way ANOVA, with Dunnett’s test for inset (F_2, 70_ = 0.8296; 5’-Del *p =* 0.7986, 3’-Del *p =* 0.339). (**k**) Cumulative probabilities and log-scaled cell averages of amplitude size across genotypes, compared by Levene’s Test for averaged distributions (5’-Del F = 0.1, df = 1, *p* = 0.9204; 3’-Del F = 0.0006, df = 1, *p* = 0.9801), and 1-way ANOVA, with Dunnett’s test for inset (F_2, 70_ = 0.08143; 5’-Del *p* = 0.9431, 3’-Del *p =* 0.9904). (**l**) Transformed p-values of Levene’s Test and (**m**) SynGO gene-set averaged log_2_FC values across pre- or post-synaptic genes. (**n**) Gene expression panel (z-scores), of canonical *NRXN1* binding partners in iGABA neurons, boxes indicate reaching genome wide significance. (**o**) Representative images of iGABA synaptic puncta traced (SYN1) onto dendrites (MAP2), and (**p**) normalized fold change of SYN1 puncta to length of MAP2 ratio (Syn:Neu), (n: Control = 33/2; 5’-Del: 36/2; 3’-Del: 40/2), compared via 1-way ANOVA, with Dunnett’s test (F_2, 101_ = 12.59, 5’-Del *p <* 0.0001, 3’-Del *p =* 0.0271). *\*p* < 0.05, *\*\*p* < 0.01, *\*\*\*p* <0.001, Wilcoxon’s rank sum test, FDR = 0.05. Data represented as mean ± sem. n reported as samples/donors | independent batches.

By contrast, synaptic transmission in iGABA neurons appeared to be enhanced. The frequency of spontaneous inhibitory post-synaptic currents (sIPSCs) increased, marked by a significant decrease in IEI in 3’-Del iGABA neurons (*p* = 1.00E-4 by Levene’s test with Bonferroni correction) (**Fig. 4i,j**). There was no change in sIPSC amplitude (**Fig. 4i,k**). Miniature inhibitory post-synaptic currents (mIPSCs) recorded in the presence of TTX and CNQX (AMPA/Kinate receptor antagonist) revealed similar trends in IEI, with no change in mIPSC amplitudes (**Extended Data Fig. 4e,f**). Similarly, SynGO analysis revealed concordant changes in synaptic transmission and transcriptomic signatures at 3’-Del (Pre-SynGO Log_2_FC = 0.0324) and represented DEGs among *NRXN1* trans-synaptic interaction partners *CBLN2*, *NLGN1*, *NXPH1, CASK,* and *LRRTM2-4* (**Fig. 4l-n**).

Quantification of synaptic puncta (via immunostaining against synapsin-1, SYN1), normalized to dendritic length (via immunostaining somatodendritic marker MAP2) uncovered a bidirectional decrease in iGLUT neurons (**Fig. 4g,h**), and an increase in iGABA neurons (**Fig. 4o,p**). Thus, divergent neurotransmission phenotypes appear to correlate with synapse number. Overall, patient-specific alterations in spontaneous neural activity were driven by synaptic deficits, with the cell type-specific impact of aberrant *NRXN1* splicing resulting in divergent neurotransmission phenotypes and changes in synapse number. Furthermore, non-recurrent LOF and GOF presented unequal effect sizes between cell-types, with 5’-Del neurons being most impacted by excitatory transmission, but 3’-Del neurons being affected by both excitatory and inhibitory transmission. Given that deletions affected iGLUT and iGABA neurons in opposing directions, these findings implicate *NRXN1* as a key mediator of excitatory/inhibitory balance, a prevalent theme among neuropsychiatric disorders^38^.

### Isogenic validation of bidirectional excitatory-inhibitory (E-I) synaptic deficits

To demonstrate a direct link between aberrant *NRXN1* splicing and synaptic dysfunction, we designed an experiment to specifically target splicing patterns. We utilized short hairpin RNAs (shRNAs) to knockdown wildtype splice isoforms in control lines, mimicking a LOF phenotype. Targeted knockdown of constitutively expressed exon 9 (expressed in alpha isoforms but not beta isoforms) achieved mRNA knockdown of ∼55% in iGLUT neurons, and ∼75% in iGABA neurons across one or more isogenic pairs (compared to a non-targeting (NT) control shRNA) (**Extended Data Fig. 6a,e**). These knockdowns would roughly mimic a 5’-Del heterozygote. Differential splicing analysis confirmed the reduction of β→α splicing in both iGLUT and iGABA neurons by ΔPSI −0.191 (*p =* 0.0083) and −0.208 (*p =* 0.0202), respectively (**Fig. 5a, i**), without significantly altering other *NRXN1* splice sites. Functionally, we observed decreased synaptic transmission in iGLUT neurons (i.e., increased sEPSC IEI, *p* = 2.2E-16 by Levene’s Test) (**Fig. 5b,c**), and increased synaptic transmission in iGABA neurons (decreased IEI, *p* = 3.65E-8 by Levene’s Test and *p* = 0.0092 by Student’s t-test) (**Fig. 5j,k**), similar to the changes with 5’-Del neurons (**Fig. 4b,j**). Transcriptomic profiles of isogenic lines further validated *NRXN1* knockdown (**Extended Data Fig 6c,g**) and demonstrated DEGs related to multiple aspects of synaptic function in both iGLUT (29/258 DEGs; 1.274161-fold, *p* = 0.1049) and iGABA neurons (384/3525 genes; 1.234861-fold, *p* = 1.307E-6) (**Fig. 5d,i and Supplementary Table S2**). Altogether, knockdown of wildtype splicing recapitulated cell-type-specific differences in *NRXN1^+/-^* 5’-Del neurons, causally implicating decreased wildtype *NRXN1α* expression (LOF) as a driver of cell-type-specific phenotypes.

**Figure 5:**
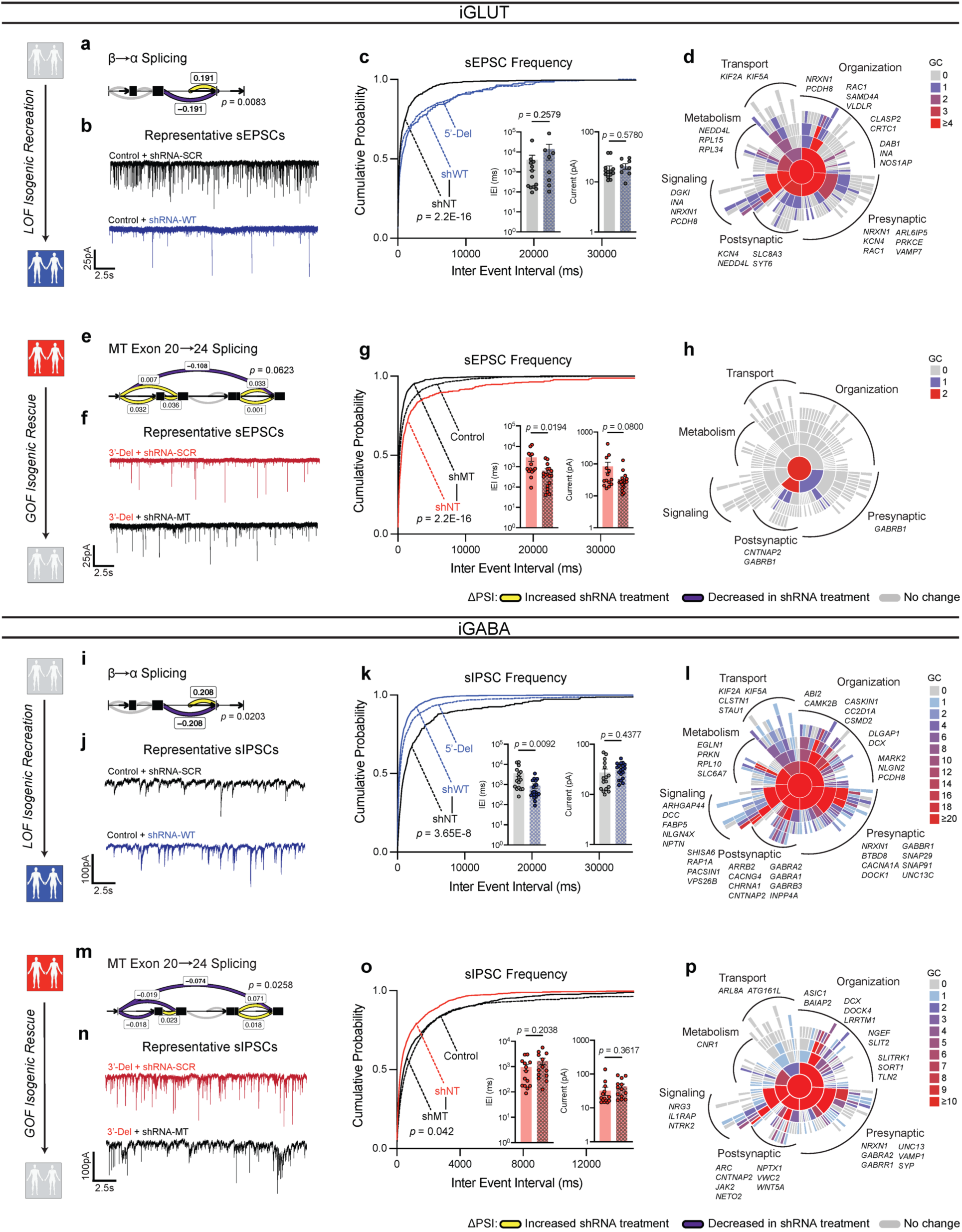
*Isogenic recapitulation and rescue of neurotransmission phenotypes.* (**a**) Differential splicing of β→α cluster in shWT, compared via Dirichlet-multinomial generalized linear model (*p* = 0.0083) in iGLUT neurons. (**b**) Representative traces of iGLUT WT knockdown effects, with (**c**) cumulative probabilities of sEPSC IEI distributions, with insets of cell-averaged IEI and amplitude measures (n: shNT = 12/1; shWT = 7/1 | 2 batches). Curves were compared via a Levene’s test (F = 70.78, df =1, *p* < 2.2E-16), and insets were compared via a Student’s t-test (IEI t=1.166, df=19, *p =* 0.2579; AMP t=0.5661, df=19, *p =* 0.578). (**d**) SynGO (biological process) sunburst plots showing enrichment of DEGs associated with synaptic function for iGLUT WT-KD compared to a brain expressed background via Fisher’s exact test (1.274161-fold, *p* = 0.1049). (**e**) Splicing of MT exon 20®24 cluster in shMT, compared via Dirichlet-multinomial generalized linear model (*p* = 0.0623). (**f**) Representative traces of iGLUT MT knockdown effects, with (**g**) cumulative probabilities of sEPSC IEI distributions, with insets of cell-averaged IEI and amplitude measures (n:shNT = 14/1; shMT = 18/1 | 2 batches). Curves were compared via a Levene’s test (F = 230.6; df = 1, *p* < 2.2E-16), and insets were compared via a Student’s t-test (IEI t=2.471, df=30, *p =* 0.0194; AMP t=1.812, df=30, *p =* 0.08). (**h**) SynGO (biological process) sunburst plots showing enrichment of DEGs associated with synaptic function for iGLUT MT-KD compared to a brain expressed background via Fisher’s exact test (1.214533-fold, *p* = 0.4541). (**i**) Differential splicing of β→α cluster in shWT, compared via Dirichlet-multinomial generalized linear model (*p* = 0.0203) in iGABA neurons. (**j**) Representative traces of iGABA WT knockdown effects, with (**k**) cumulative probabilities of sIPSC IEI distributions, with insets of cell-averaged, IEI and amplitude measures (shNT = 16/3; shWT = 19/3 | 2 batches). Curves were compared via a Levene’s test (F = 30.879, df =1, *p* < 3.66E-08), and insets were compared via a Student’s t-test (IEI t=2.768, df=33, *p =* 0.0092; AMP t=0.7856, df=33, *p =* 0.4377). (**l**) SynGO (biological process) sunburst plots for iGABA WT-KD compared to a brain expressed background via Fisher’s exact test (1.234861-fold, *p* = 1.307E-6). (**m**) Splicing of MT exon 20®24 cluster in shMT, compared via Dirichlet-multinomial generalized linear model (*p* = 0.0258) in iGABA neurons. (**n**) Representative traces of iGABA MT knockdown effects, with (**o**) cumulative probabilities of sIPSC IEI distributions, with insets of cell-averaged, IEI and amplitude measures (shNT = 16/1; shMT = 19/1 | 2 batches). Curves were compared via a Levene’s test (F = 4.1324; df = 1, *p* < 0.04226), and insets were compared via a Student’s t-test (IEI t=1.305, df=25, *p =* 0.2038; AMP t=0.9292, df=25, *p =* 0.3617). (**p**) SynGO (biological process) sunburst plots for iGABA WT-KD compared to a brain expressed background via Fisher’s exact test (1.347225-fold, *p* = 2.694E-4). Data represented as mean ± sem. n reported as samples/donors | independent batches.

For GOF *NRXN1 3*’-Del neurons, we took a different approach and applied shRNAs to knockdown mutant splice isoforms, in an attempt to reverse the GOF phenotype. We designed a shRNA against the mutant splice junction overlapping exons 20 and 24, expressed in all 3’-Del unique *NRXN1* alpha and beta isoforms, and achieved targeted knockdown of mutant splice isoforms by 95% in iGLUT, and 25% in iGABA neurons in both donors (**Extended Data Fig 6b,f**). Differential splicing analysis again confirmed the selective reduction of GOF splicing in both iGLUT and iGABA neurons, by ΔPSI −0.108 (*p =* 0.0623) and −0.074 (*p =* 0.0258), respectively (**Fig**. **5e,m**), without significantly altering other *NRXN1* splice sites. In the donor with the most robust knockdown in both iGLUT (90%) and iGABA (37%) neurons, electrophysiological recordings revealed a reversal of synaptic transmission phenotypes, achieving IEI levels similar to control iGLUT and iGABA neurons (**Fig. 4b,j**). In iGLUT neurons, shRNA MT decreased sEPSC IEI (*p* < 2.2E-16 by Levene’s Test and *p* = 0.0194) (**Fig. 5f,g**) and in iGABA neurons increased sIPSC IEI (*p* = 0.042 by Levene’s Test) (**Fig 5h,o**), as compared to shRNA-NT. Parallel to the LOF isogenic transcriptomic profiles, the GOF isogenic line had modest changes in iGLUT neurons (3/28 DEGs; 1.214533-fold, *p* = 0.4541 by Levene’s test), but more pronounced synaptic DEGs in iGABA neurons (128/1077 DEGs; 1.347225-fold, *p* = 2.694E-4) (**Fig. 5h,p and Extended Data Fig. 6d,h**).

In summary, knockdown of WT *NRXN1* in iGLUT and iGABA neurons recapitulated the consequences of reduced β→α splicing on neurotransmitter phenotypes of 5’ LOF *NRXN1* deletions, while knockdown of MT splicing appeared to mitigate the negative effect of the 3’ GOF *NRXN1* deletions. Furthermore, the bidirectional effect of *NRXN1* deletions on iGLUTs (decrease) and iGABAs (increase), i.e., E-I balance, was reversed by shRNA treatments. Transcriptomic profiling of shRNA treated isogenic lines revealed ∼10% of DEGs related to synaptic function across all four experimental conditions. Altogether, shRNA mediated perturbations among WT and MT isoforms causally implicate *NRXN1* splicing to synaptic dysfunction.

### Framework for precision medicine against stratified LOF/GOF phenotypes

As a proof-of-principle therapeutic intervention, we tested methods to directly and indirectly manipulate *NRXN1* expression, separately targeting LOF and GOF mechanisms, and focusing on reversing decreased excitation phenotypes in iGLUT neurons.

For LOF patients, we hypothesized that increasing transcription of the WT *NRXN1α* allele would restore *NRXN1* levels and reverse the signature of reduced β→α splicing (**Fig. 6a**). β-estradiol reversed *NRXN1* LOF neurogenesis deficits in *xenopus* and human NPC models^39^. Although the mechanism is unknown, chromatin immunoprecipitation with sequencing (ChIP-Seq) in mouse brain tissue identified estrogen receptor alphaα (ERα) binding sites at the *NRXN1* alpha locus (**Extended Data Fig. 7a,b**)^40^. We predicted *NRXN1* to be a target of ERα, and report that acute treatment with β-estradiol (10nM or 30nM, 3-5 days) significantly increased *NRXN1α* expression in iGLUT neurons (*p* = 0.0297 by Student’s t-test) derived from 5’-Del patients (**Fig. 6b**), but not in controls (**Extended Data Fig. 7c**). The functional effect of chronic treatment of post-mitotic 5’-Del neurons with 30nM β-estradiol (relative to DMSO vehicle) was evaluated across spontaneous neural activity (MEA), synaptic transmission (patch-clamp) and gene expression (RNA-sequencing). MEA recordings at WPI3 revealed a significant increase in wMFR activity (*p* = 0.0057 via student’s t-test) in the β-estradiol treated condition (**Fig. 6c**). Transcriptional profiling of the β-estradiol treatment detected an inversion of the β→α splicing signature (ΔPSI 0.045), and more subtle changes in global gene expression (7 upregulated and 34 downregulated DEGs) (**Fig. 6d,e**). Likewise, patch-clamp recordings revealed that β-estradiol treatment ameliorated the sEPSC IEI phenotype in 5’-Del neurons (*p =* 4.62E-4 by Levene’s Test, Bonferroni corrected), but not in vehicle treated 5’-Del neurons (*p =* 0.9055 by Levene’s Test), compared to healthy controls (**Fig. 6f,g**).

**Figure 6:**
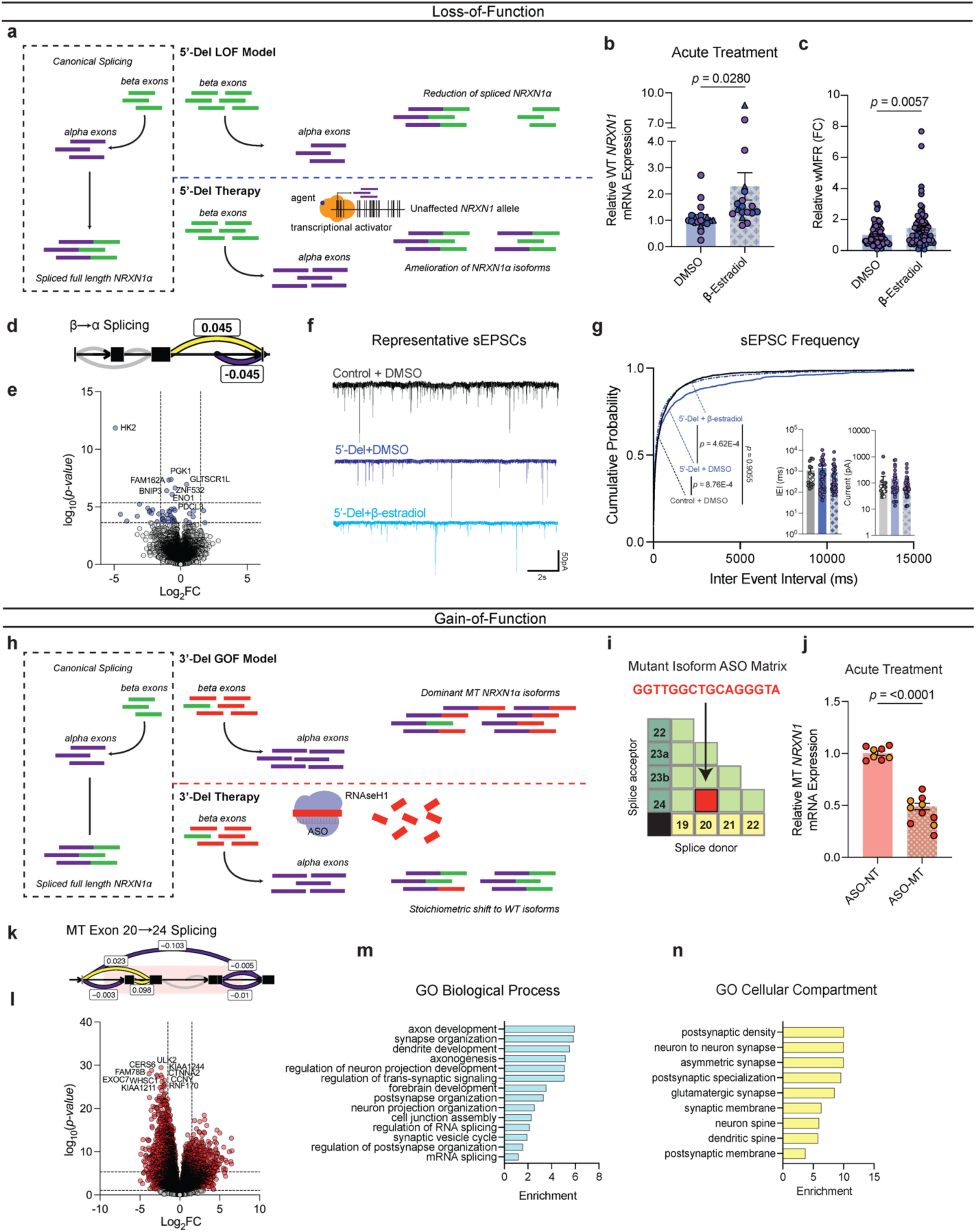
*Precise therapeutic targeting of stratified GOF- and LOF-NRXN1^+/-^ in iGLUT neurons.* (**a**) Model and proposed mechanism of rescue for LOF patients ameliorating loss of wildtype isoforms. (**b**) RT-qPCR of WT *NRXN1* gene expression in 5-Del patients, post-acute treatment (3-5 days) (n: DMSO = 19/2; β-estradiol = 18/2 | 3 batches) compared via Student’s t-test t=2.293, df=35, *p =* 0.028. (**c**) Quantification of iGLUT neuronal activity at WPI3 across vehicle and treatment (n: DMSO = 72/3; β-estradiol = 74/3 | 3 batches) compared via Student’s t-test (t=2.804, df=144, *p* = 0.0057). (**d**) Splicing of β→α cluster and (**e**) volcano plot of DEGs in β-estradiol treated 5’-Del iGLUT neurons, compared to vehicle treated 5’-Del iGLUT neurons. (**f**) Representative patch-clamp traces and (**h**) cumulative probabilities of sEPSC IEI distributions, with insets of cell-averaged, IEI and amplitude measures (n: Control + DMSO = 18/1; 5’-Del + DMSO = 33/2 5’-Del + β-estradiol = 35/2). Curves were compared via Levene’s test with Bonferroni corrections (Control + DMSO v. 5’-Del + DMSO F = 13.151; df = 1, *p* = 8.763E-4; 5’-Del + DMSO v 5’-Del + β-estradiol F = 14.359; df = 1, *p* = 4.62E-04; Control + DMSO v 5’-Del + β-estradiol, F = 0.0141; df = 1, *p* = 0.955). Insets were compared via Student’s t-test (IEI t=1.161, df=66, *p =* 0.2499). (**h**) Model and proposed mechanism of rescue for GOF patients expressing mutant isoforms. (**i**) Schematic of a *NRXN1* ASO matrix, with the selective sequence targeting 20/24 splice junction. (**j**) RT-qPCR of MT *NRXN1* gene expression in 3’-Del patients’ post-acute treatment for 72hrs (ASO-NT = 8/2; ASO-MT = 10/2 | batches),compared via Student’s t-test t=12.91, df=16 *p <* 0.0001. (**k**) Splicing of MT Exon 20→24 cluster and (**l**) volcano plot of DEGs in ASO-MT treated 3’-Del iGLUT neurons, compared to ASO-NT treated 3’-Del iGLUT neurons.(**m**) Biological process and (**n**) cellular compartment GO terms demonstrating enrichment (-log_10_[adj *p* value]) of selected synapse related pathways. Data represented as mean ± sem. n reported as samples/donors | independent batches.

Towards a therapeutic strategy for treating GOF patients, we investigated the utility of anti-sense oligonucleotides (ASOs), recently used to treat several neurological diseases^41^, to target a specific RNA splice site. We designed an “alternative splice matrix” by juxtaposing splice donor and acceptor RNA sequences from each *NRXN1α* exon, generating all possible combinations of canonical and non-canonical splice junctions (**Fig. 6h,i**). Then, for the sequence covering the 20/24 mutant splice junction, we designed an ASO to facilitate targeted RNAseH1-dependent degradation of 3-Del mutant isoforms. ASO treatment (1μM) for 72hrs decreased total mutant isoforms by ∼55% in iGLUT neurons (*p* < 0.001 by student’s t-test), as compared to a non-targeting control ASO (**Fig. 6j**). Differential splicing analysis confirmed the reduction of GOF splicing (ΔPSI −0.103) and increased ratios of SS4+. We also observed a significant reduction in β→α splicing (ΔPSI −0.253, *p =* 0.007, Bonferroni corrected), perhaps resulting from reduced MT isoform expression, contrasting the elevated baseline of *NRXN1α* in 3’-Del (**Fig. 6k**). Transcriptomic profiling revealed robust DEGs, enriched for synaptic properties, neurotransmitter signaling, and neurodevelopmental pathways **(Supplementary Table S3)** (231/1844 Bonferroni corrected DEGs annotated in SynGO; 1.420028-fold, *p* = 1.016E-08), (**Fig. 6c-d**). To demonstrate the broader applicability of an ASO strategy, we report mutant splice isoforms in post-mortem brain tissue from an unrelated *NRXN1^+/-^* case diagnosed with autism spectrum disorder (ASD) (**Supplemental Fig. 8**). By integrating long-read and short-read sequencing^15^, we identified nine high-confidence isoforms that are predicted to be translated: two of which contained a novel splice junction overlapping the deletion encompassing exon 14/19, targetable by an ASO tailored to the mutant splice junction.

Altogether, increased *NRXN1* expression in LOF neurons and knockdown of mutant splicing in GOF neurons can each rescue the splicing defects that produce opposing cell-type-specific case/control differences in *NRXN1^+/-^* neurons. We propose the aberrant *NRXN1α* splicing is a key a driver of complex *NRXN1^+/-^*phenotypes and that modulators of *NRXN1* expression and/or splicing represent novel targeted therapies.

## DISCUSSION

*NRXN1^+/-^* deletions are non-recurrent between patients, linked to diverse clinical outcomes that cannot be explained by the size or boundaries of the deletion itself^6^. Here, we show that *NRXN1* splicing alters excitatory-inhibitory (E-I) imbalance^1,32^, a common theme among neuropsychiatric disorders, by bidirectionally regulating synaptic transmission, with a decrease in frequency of sEPSCs in 5’-Del and 3’-Del iGLUT neurons but an increase in frequency of sIPSCs in 3’-Del iGABA neurons. Using a case/control *NRXN1^+/-^* cohort as well as isogenic manipulations of the *NRXN1* isoform repertoire, we report distinct phenotypic effects in human iGLUT and iGABA neurons that predominately manifest in changes in the frequency of synaptic function. These results suggest causal relationships between aberrant *NRXN1* splicing and synaptic dysfunction, dictated by unique patient-specific mechanisms.

The regulation of *NRXN1* splicing, including the formation of aberrant splice sites by specific RBPs, is poorly understood; >100 RBPs are predicted to interact with *NRXN1* mRNA in a cell-type^42,43^ and neuronal activity^23^ dependent-manner. For example, KH-domain STAR-family RBPs regulate SS4+ in a neuronal activity dependent manner^23–25^, mediating trans-synaptic signaling by varying interactions among a host of post-synaptic ligands. Although patient transcriptomic profiles nominate STAR proteins as potential drivers of aberrant GOF splicing, further investigation is required to identify the biochemical mechanisms involved. Nevertheless, we posit that direct manipulation of splicing may achieve therapeutic benefit in some cases.

Neurexins are expressed across all synapses and among certain non-neuronal cell-types, such as astrocytes, perhaps with distinct cell-type-specific functional roles^37^. Mechanistically, our study did not resolve whether mutant GOF isoforms shifted the stoichiometry of alternative splicing against wildtype RNA isoforms or if they altered trans-synaptic protein-protein interactions. In either case, *NRXN1* encodes a pre-synaptic molecule but traditional neuropharmacological agents typically target specific receptors. Given that patient-specific patterns of aberrant *NRXN1* splicing across non-recurrent mutations are unlikely to be reversed by a pharmacological approach, precision therapies that target aberrant splicing are needed to effectively reverse defects in the multiple cell types affected by patient-specific deletions.

The opposing effects of *NRXN1^+/-^* in glutamate and GABA neurons provides the foundation for evaluating proof-of-concept therapeutics that target distinct LOF or GOF mechanisms. For LOF mutations, mechanisms for upregulation of wild-type allelic expression and/or restoration of proper *NRXN1* splicing are required (**Fig. 6a**). While direct approaches to increase *NRXN1* expression may be ideal, such as small activating RNAs^44^, further work is required to carefully evaluate safety and efficacy in clinical trials. On the other hand, indirect approaches have the advantage of prioritizing drugs from a list of already clinically approved molecules, selected based on their predicted ability to alter the expression of a target gene (e.g., LINCs^45^). With relevance to the ability of estradiol to increase *NRXN1* and rescue synaptic deficits, steroid-based pharmaceuticals are already used to treat neuroinflammation-related conditions^46^, with recent studies revealing estrogen-mediated roles in neuroprotection^39^, demonstrating the safety and feasibility of this approach. For GOF, ASOs designed to target a mutant splice junction and facilitate degradation of MT isoforms may represent a viable therapeutic approach (**Fig. 6h**). Not all in-frame coding mutations will be GOF; for example, unique mutations at non-canonical splice donor/acceptor sites will require further characterization prior to assigning LOF/GOF status. In all cases, maintaining a stoichiometric balance between MT and WT isoforms may require combination treatments (e.g., ASO and transcriptional activator) for therapeutic benefit. Cases of muscular dystrophies treated via splicing modulating ASOs have proven effective^47^, but overall, gene therapy in brain disease has shown mixed successes^48^. Critically, although our proof-of-principle *in vitro* findings suggest a novel therapeutic avenue, translational studies will be required to confirm the efficacy of this framework *in vivo*, and ultimately, in the clinic.

Several technical limitations warrant acknowledgement. The iGLUT and iGABA neuron populations studied herein are not just immature, but also comprise diverse neuronal subtypes^49^; thus, the more discrete impacts of *NRXN1^+/-^* on synaptic physiology across different subtypes of neurons (e.g., SST versus PV expressing GABA neurons) remain unresolved. Moreover, future studies in more physiologically relevant and circuit-like models may uncover novel non-autonomous and activity-dependent phenotypes. Mechanistically, it will be important to probe the biochemical and proteomic interactions of *NRXN1*; for example, unbiased proximity-labelling methods (e.g., BioID^50^) could define perturbations in protein-protein interaction profiles between wildtype and mutant isoforms.

Precision medicine seeks to tailor treatments to individual patients,^51^ regardless of differential penetrance of genetic mutations. Therefore, we focused on stratifying mutations in the same gene based on LOF and GOF mechanisms, in contrast to clinical diagnosis (bipolar disorder or schizophrenia), with the goal of one day providing the right therapy at the right time to the right patient. The most successful examples of precision medicine to date have been in cancer^52^ and monogenic disease^53^, whereby genetic analyses revealed molecular subtypes that benefited from specific treatments. In the context of brain disorders, patient stratification is particularly challenging, reflecting in part the myriad rare and common variants linked to disease^54^. Disease-agnostic analyses reveal that transcription factors and nucleic acid binding proteins are overwhelmingly driven by LOF mutations, whereas signaling molecules, enzymes, receptors and transporters more frequently incur dominant GOF mutations^55^. When both are possible, distinct mutations in the same genes can result in pleiotropic phenotypic effects. A binary therapeutic approach, similar to what we proposed here for *NRXN1*, may prove suitable for dual LOF an GOF mechanisms linked to mutations in other neuropsychiatric disorder-related synaptic genes (e.g., *NLGN3*^56^, *CACNA1D*^57^, *CACNA1C*^58^*, SCN1A*^59^*, SCN2A*^60^) as well as broadly in neurodegenerative disease (e.g., *SOD1*^61^*, TDP43*^61^*, FUS/TLS*^61^*, C9ORF72*^62^, *AR*^63^), and many short tandem repeat disorders^64^. Taken together, our work advances precision medicine, demonstrating the necessity of functionally dissecting the phenotypic impact of diverse patient-specific genetic variants across cellular contexts, in order to resolve candidate therapies across stratified disease mechanisms.

## METHODS

### Plasmid designs and molecular cloning

#### i. TetO-Ascl1-Neo

The *ASCL1* insert from TetO-Ascl1-Puro (Addgene #97329) was synthesized as a gBLOCK flanked by EcoR1 cut sites, and cloned into TetO-hNgn2-Neo using EcoR1 to remove *NGN2.* The recipient vector was dephosphorylated with shrimp alkaline phosphatase (rSAP NEB #M0371S) during the digest, column purified and ligated at a 1:1 vector to insert ratio using the QuickLig Kit (NEB #M2200S). The *ASCL1* stop codon was subsequently mutated using the QuickChange II-XL site-directed mutagenesis kit (Agilent #200523) and verified via whole plasmid sequencing from plasmidsaurus.

#### ii. shRNA RNA interference constructs

All shRNAs were designed and produced by Sigma-Aldrich via custom submitted sequences against wildtype *NRXN1ɑ* (constitutively expressed exon 9) and mutant *NRXN1ɑ* (mutant 20/24 splice junction) cloned into TCR2 pLKO.5-puro.

### Cell Culture

#### i. hiPSC culture

Passage matched (±3) human induced pluripotent stem cells (hiPSCs) were cultured in StemFlex media (Life technologies #A3349401) on Matrigel (Corning, #354230). At ∼70-80% hiPSCs were clump passaged using 0.5mM EDTA in PBS without Mg/Ca (Life technologies #10010-031). Cells were washed once and incubated for 4-7 min with 0.5µM EDTA at RT. The EDTA was aspirated, and the remaining colonies were lifted off with 1mL of StemFlex and re-distributed at varying densities in a 6–well maintenance plate. hiPSC lines were split every 4-6 days. For neuronal/organoid differentiation, wells of similar confluence across all hiPSC donors were resuspended and seeded onto a Matrigel coated 10cm dish and expanded until ∼70-80% confluency.

#### ii. HEK293T culture and lenti-viral production

HEK293T cells were maintained in 15cm dishes and cultured in DMEM supplemented with 10% standard FBS (Life technologies #16000069). 3rd Gen lenti-viral particles were produced using previously described methods and REV, MDL and VSV-G plasmid ratios^18^, each transfected with PEImax (Polysciences #24765-1). Each PEIMax batch was volumetrically titrated at total µgDNA: µLPEI for optimal transfection efficiency.

#### iii. Primary mouse glia production

All mouse work was performed under approved IACUC protocols at the Icahn School of Medicine at Mount Sinai. C57BL/6 mice were used as breeding pairs. For glial preps, dissected cortices from 3 pups (at p0-3) were dissociated using papain (R&D #LS003126) and plated on 10cm dishes in MEF medium (DMEM, 10% Cosmic Calf Serum (Fisher #SH3008703HI), 1x Antibiotic-antimycotic (Life technologies #15240), 1x Sodium Pyruvate (Life technologies #11360070), 1x MEM Non-Essential Amino Acids Solution (Life technologies #11140050), 4uL 2-Mercaptoethanol (Sigma #60-24-2), supplemented with 100µg/mL Normocin (InvivoGen #ant-nr-2). Glial cells were recovered and propagated for 7 days, and expanded into three 10cm dishes. To promote neuronal health and synapse maturation, we utilized mouse glia from well-established protocols that significantly outperform human astrocytes for co-culture experiments^19,21,65^. All glial preps were tested twice for mycoplasma (Normocin withdrawn) (Lonza, #LT07-318) prior to freezing or neuronal co-culture. At day 14, one 10cm dish with mouse glia were distributed to two MEA, 12- or 24-well plates, and subsequently inactivated with 4µM Ara-C (Sigma #C1768) prior to or during re-seeding of induced neurons.

#### iv. iGLUT induction and astrocyte co-culture

At day −1 hiPSCs expanded in 10cm dishes were dissociated with accutase (StemCell Technologies, #07920), washed and pelleted with a 1:4 ratio of accutase to DMEM for 5 min at 1000 rcf, and re-suspended with StemFlex media containing ROCK inhibitor, THX (10µM/mL; Sigma Aldrich, SML1045). The hiPSCs are then co-transduced with TetO-Ngn2-Puro (Addgene #79049) or TetO-Ngn2-Neo (Addgene# 99378) and UCb-rtTA (legacy vector from the lab of Fred Gage) and seeded at 1.0-1.5×10^6^ cells in 1.5mL per well on 6-well plates were coated with 2x Matrigel for at least one hour at 37°C. The hiPSC-viral mixture was then incubated overnight. The following morning (day 0), a full media change with iGLUT induction media was performed with the following recipe: Neurobasal Media: 1x N2 (Life technologies #17502-048), 1x B-27 minus Vitamin A (Life technologies #12587-010), 1x Antibiotic-Antimycotic, 1x Sodium Pyruvate, 1x GlutaMax (Life technologies #35050), 500µg/mL cyclic-AMP (Sigma #D0627), 200µM L-ascorbic acid (Sigma #A0278), 20ng/ml BDNF (Peprotech #450-02), 20ng/ml GDNF (Peprotech #450-10), 1µg/ml natural mouse laminin (Life technologies #23017015). On days 1-2, iGLUT cells were treated with respective antibiotic concentrations at 1µg/mL puromycin (Sigma# P7255) or 0.5µg/mL neomycin (Life technologies #11811-031). On day 3, antibiotic medium was withdrawn and iGLUT cells were treated with 4µM Ara-C. On Day 4, iGLUT cells were dissociated with accutase for 15 min, washed and pelleted with a 1:4 ratio of accutase to DMEM for 5 min at 800 rcf, and re-suspended with iGLUT media containing ROCK inhibitor, Ara-C and 2% low-hemoglobin FBS (R&D systems #S11510). iGLUT neurons were distributed among wells (500-750k cells per 24wp or 0.75-1.5E6 cell per 12wp) pre-seeded with confluent mouse glia. The following day, iGLUT neurons received a full media change with Brainphys maturation media (Neurobasal Media, 1x N2, 1x B-27 minus Vitamin A, 1x Antibiotic-Antimycotic, 500µg/mL cyclic-AMP, 200µM Ascorbic Acid, 20ng/ml BDNF, 20ng/ml GDNF, 2% low-hemoglobin FBS, 1µg/ml Mouse Laminin) supplemented with Ara-C and were subsequently monitored for growth of non-neuronal/glial cells. Ara-C treatment was tittered down with half-media changes (without Ara-C) every 3-4 days until used for experiments.

#### v. iGABA induction and astrocyte co-culture

iGABA production paralleled the methods aforementioned. hiPSCs were instead co-transduced with TetO-Ascl1-puro (Addgene #97330) or TetO-Ascl1-neo (Addgene #TBD), TetO-Dlx2-hygro (Addgene #97329) and UCb-rtTA and seeded at 0.8-1.2×10^6^ cells in 1.5mL per well on 6-well plates similarly prepared. The following morning (day 0), a full media change with iGABA induction media (DMEM/F-12 + Glutamax, 1x N2, 1x B-27 minus Vitamin A, 1x Antibiotic-Antimycotic) was performed. On days 1-2 iGABA cells were selected with respective antibiotic concentrations at 1µg/mL puromycin or 0.5µg/mL neomycin and 0.25µg/mL hygromycin (Life Technologies #10687010), followed by antibiotic withdrawal and Ara-C treatment on day 3. iGABA neurons were re-seeded identically to iGLUT cells, at 150-250k cells per 24wp well. iGABA cultures were morphologically QC’ed prior to all experiments, with uncharacteristic batches being discarded.

#### vi. Cortical and subpallial organoid differentiation

Cortical organoids were generated according to the protocol described by Sloan et. al.^66^, with several modifications. hiPSCs were first aggregated into embryoid bodies (EBs) using an AggreWell™800 Microwell Culture Plate system (Stemcell Tech #34850). Expanded hiPSCs were rinsed twice with DPBS without Ca/Mg, and then dissociated using accutase. 3×10E6 hiPSCs were added to a single well in the prepared AggreWell and allowed to aggregate in a 37°C incubator for 24 hours. The following day (day 0), EBs were dislodged from the AggreWell plate using a cut p1000 pipette tip and passed over a 40μm strainer, and washed with excess DMEM. The strainer was inverted over an Ultra-Low Attachment 10 cm culture dish and the EBs were collected in spheroid induction media, which contained Stemflex supplemented with two SMAD inhibitors, SB-431542 (SB) and LDN193189 (LDN), and THX. The following day (day 1), the media THX was withdrawn. From d2-d6, induction media was replaced daily, and no longer contained the Stemflex supplement (only base Stemflex media with SB and LDN). On day 6, the media was replaced with organoid patterning media, formulated with Neurobasal-A medium, 1x B-27 minus Vitamin A, 1x GlutaMAX, and 1x Antibiotic-Antimycotic. From d6-d24, organoid maturation media was supplemented with 20ng/ml of EGF (R&D Systems, #236-EG) and 20ng/ml FGF2 (R&D Systems, #233-FB-01M). Media was changed every day from d6 – d15, after which media was changed every other day. From d25-d43, the organoid maturation media was supplemented with 20ng/ml of NT-3 (PeproTech, #450-03) and 20ng/ml BDNF. From d43 onwards, organoids received organoid maturation media with no supplements, and media was changed every 4 days or as needed. Subpallial organoids were generated in the same way as cortical organoids, but with additions in media formulations. From d4-d23, hSOs received spheroid induction media or organoid maturation media supplemented with 5µM of the Wnt inhibitor, IWP-2 (Selleckchem, #S7085). From d12-d23, hSO organoids received neuronal differentiation media supplemented with 100nM of the SHH agonist, SAG (Selleckchem, #S7779).

### Electrophysiology

#### i. Multi-electrode array (MEA)

The Axion Maestro (Middleman) system was used to perform all MEA recordings. Following iGLUT and iGABA inductions, 80-100k cells were re-plated on each MEA well and measurements began as early as DIV9 for both iGLUT and iGABA co-cultures. For time course experiments, MEA plates were recorded every 2-3 days per week with a full media change prior to each recording. Comparisons for iGLUT neurons were made at WPI4 and WPI6, as well characterized timepoints for synaptic activity. For iGABA neurons, comparisons were made at WPI2 and WPI6 to include both timepoints of elevated neuronal activity. For acute drug treatments with gabazine (Tocris, #1262), a baseline recording (pre-treatment) was first obtained, followed by an immediate addition of a small volume of concentrated stock. A second, post-treatment recording was then obtained to evaluate the difference in activity before and after drug treatment. A full media change was performed one day prior to the day of recording. MEA wells were visually QC’ed for similar densities prior to recording, with high/low density wells being excluded from recordings.

#### ii. Whole-cell patch-clamp electrophysiology

For whole-cell patch-clamp recordings, iGLUT (300k/well) or iGABA (250k/well) human-mouse glia co-cultures were recorded at 4-6 weeks following dox-induction (time points specified in figure legends), with a full media change one day prior to recording. Only coverslips with similar densities were selected for recording. Cells were visualized on a Nikon inverted microscope equipped with fluorescence and Hoffman optics. Neurons were recorded with an Axopatch 200B amplifier (Molecular Devices), digitized at 10 kHz using a Digidata 1320a (Molecular Devices) and filtered between 1-10 kHz, using Clampex 10 software (Molecular Devices). Patch pipettes were pulled from borosilicate glass electrodes (Warner Instruments) to a final tip resistance of 3-5 MΩ using a vertical gravity puller (Narishige). To sustain the baseline activity of neurons from extended cultures, coverslips were recorded in base Brainphys medium (external solution, Osm 305, pH 7.49). Each coverslip was equilibrated to room temperature for 10min, with 3-5 neurons were recorded for no more than a total of 75min per coverslip. For measurements of passive and excitable properties, an internal patch solution was used containing (in mM): K-d-gluconate, 140; NaCl, 4; MgCl2, 2; EGTA, 1.1; HEPES, 5; Na2ATP, 2; sodium creatine phosphate, 5; Na3GTP, 0.6, at a pH of 7.4. Osmolarity was 290-295 mOsm. Neurons were chosen at random using DIC and all recordings were made at room temperature (∼22°C). Current-clamp measurement occurred across −10pA to +50pA steps, with a maximum stimulus of +60pA, whereas voltage-clamp measurements occurred across −50mV to +50mV steps, normalized to cell capacitance (to control for variable neuronal size). Current clamp measurements were corrected for the junction potential (∼-15.5mV). For sEPSC/sIPSC recordings, the internal solution was replaced with (in mM): Cesium-Chloride, 135; HEPES-CoOH, 10; QX-314, 5; EGTA, 5. Osmolarity was 290-295 mOsm. All mEPSC measurements were recorded under the presence of 100nM TTX-citrate (Tocris Cat# 1069). mIPSC measurements were made using 100nM TTX-citrate and CNQX-disodium salt (Tocris Cat#1045/1) to pharmacologically inhibit ionotropic glutamate receptors. All chemicals were purchased from Sigma-Aldrich Co. (St. Louis, MO). All toxic compounds were handled in accordance with ISMMS EHS standards.

#### iii. Patch-clamp data analysis

All patch-clamp data were analyzed on Clampfit (v11) and Easy-Electrophysiology (v2.4.1 or beta-versions). Briefly, for voltage-clamp data, files were opened in Easy-Electrophysiology and two bins were assigned for Na+/K+ measures for minimum and maximum current values, respectively. For current-clamp data, files were opened in Easy-Electrophysiology and action potential (AP) analysis was used to automatically determine spike number and properties. For gap-free recordings, all data was post-hoc adjusted to a baseline of zero on ClampFit, and subsequently analyzed in Easy-Electrophysiology by template and linear threshold detection. For case v. control experiments, a minimum cutoff of 10 events for the duration of the recording (3min) was used as QC. For typical EPSC events, a single template from a randomly chosen recording was used to analyze all traces (with a 30ms decay cutoff). For IPSC events, three templates were used to detect variable GABA receptor kinetics, for all traces (with a 60ms decay cutoff). An amplitude cut off of 7pA was used to call positive events. For cumulative probabilities, each cell was binned by experiment and averaged across all cells for a representative curve (GraphPad Prism).

### RNA-Sequencing and bioinformatic analyses

#### i. Bulk RNA sequencing and DEG analysis of iGLUT and iGABA neurons

iGLUT (DIV21) and iGABA (DIV14 or DIV35) co-cultured with primary mouse glia (to match functional experiments) were harvested in Trizol and submitted to the New York Genome Center for high-throughput RNA extraction and quality control, library prep using the Kapa Hyper library prep with ribo-erase (Roche #KK8541) and subsequently sequenced on Illumina NovaSeq. Similarly, shRNA samples were harvested at DIV21-24, and DIV35-49 for iGLUT and iGABA neurons, respectively. Returned raw data files were first processed to remove mouse reads in the RNA-seq data, a combined genome reference (hg38 and mm10) was created using the “mkref” command in cellranger (v6.1.2). The raw sequencing reads were then aligned to the combined genome reference using STAR (v2.7.2a)^67^. Reads that mapped specifically to the human reference genome were extracted from the resulting BAM files for subsequent gene expression analysis. Gene-level read counts were obtained using the Subread (v2.0.1) package featureCount^68^, and RPKM values were calculated using the R package edgeR^69^. To confirm sample identity, variants were called from RNA-seq bam files by HaplotypeCaller and GenotypeGVCFs in GATK (v4.2.0). Then bcftools (v1.15) was used to examine variants concordance with variants from whole-exome sequencing data from the same donor. Following donor identity confirmation, the differential gene expression analysis followed the methods as described previously^70^. First, CibersortX^71^ was utilized to predict differences in cell type composition across all samples. The R package variancePartition (v1.30.2)^72^ was then employed to investigate the contribution of specific variables to the variance in gene expression. The limma/voom package^27^ was used for differential expression analysis, with the regression of fibroblast and hiPSC cell type compositions. The analysis began with a comparison between the case and control groups. Subsequently, within each case vs control group, subgroup comparisons were conducted for all four pairs (two donors each for 3’-Del and 5-Del patients and two healthy controls) of samples, accounting for heterogeneity between different donors. Genes with an FDR less than 0.1, and a fold change above 1.5 and below −1.5 in all four pairs of subgroup comparisons were defined as the final set of differentially expressed genes, unless otherwise specified. Kallisto (v0.46.1)^73^ was used to calculate the *NRXN1* exon usage ratios.

#### ii. Analysis of alternative splicing estimates via LeafCutter

Reads were aligned to a combined GRCh38 human and GRCm38 mouse reference genome using STAR (v2.7.1.a), with an index built against GRCh38 Gencode GTF (v92) using the option −5sjdbOverhang 100. To allow the discovery of novel splice junctions and increase mapping accuracy, STAR was run in two-pass mode with standard options. Reads mapping exclusively to GRCh38 were subsequently extracted and replicates merged with samtools (v1.6). Splice junctions were extracted from the resulting bam files with regtools v0.5.2 ‘junctions extract’ with parameters “-a 8”, “-m 50”, and “-M 1200000” due to potential for the genomic deletions to cause larger intervals between junctions. Junctions were clustered using the leafcutter pipeline (v2c9907e) script “leafcutter_cluster_regtools.py” with option “-l 1200000”. For differential splicing analysis, leafcutter was run with “-i 2 −5g 0” to reflect the samples used. Splicegraph visualizations were constructed using a modified version of the leafviz pipeline. All statistical tests utilized a Dirichlet-multinomial generalized linear model, and were corrected for multiple comparisons when necessary via Bonferroni adjustment.

#### iii. Pathway and Network Analysis of DEGs

For hierarchical clustering of DEGs based on gene expression fold change, the Pheatmap function in R was used cluster gene expression fold change of DEGs combined from both 5’-Del and 3’-Del conditions. R package clusterProfiler^29^ was used to performed the gene set enrichment analysis (GSEA). For each cell type, ranked gene expression of each genotype (5’-Del or 3’-Del) against control were used as background and DEGs from both genotypes (5’-Del and 3’-Del) were used as query gene sets. For Gene Ontology (GO) enrichment analysis, the enrichGO function within R package clusterProfiler was used. For GO with a default background, query gene list was converted from ‘ENSEMBL’ format to ‘ENTREZID’ format by bitr function in clusterProfiler and the OrgDb parameter was set as ‘org.Hs.eg.db’. When customized genes were used as background, both background and query gene lists were kept as ‘ENSEMBL’ format. For SynapseGO (SynGO v1.2), DEG or prioritized DEG lists (Log_2_FC filtered) gene lists were tested for enrichment with “brain expressed” background set selected, containing 18035 unique genes in total of which 1591 overlap with SynGO annotated genes. Sunburst plots were exported via the web-based application. The ASD, BP, and SCZ risk gene lists were extracted from previously curated gene lists^30^, Genes with the top 200 smallest FDR values and a fold change larger than 1.5 in the case vs control comparison were selected for the protein interaction network analysis. Then, overlapping genes between the selected gene list, the disease risk gene sets, and the proteins in the SIGNOR database were utilized to query the SIGNOR database and build the interaction network, using the “connect + add bridge proteins” search mode in The SIGNOR Cytoscape App (v1.2)^74^.

#### v. Dissociation and 10x Single-Cell RNA sequencing of organoids

Whole organoids were dissociated to the single cell level in preparation for single-cell RNA sequencing using the Papain Worthington Kit (Worthington, LK003150). All solutions included in the kit were prepared according to the manufacturer’s instructions. 4-6 organoids were transferred to one well of a low attachment 6 well plate and were washed with PBS without Ca^2+^ and Mg^2+^. Organoids were cut with a small scalpel into smaller pieces for easier dissociation. 800µl of papain solution (supplied in the kit) was added per well. Samples were incubated at 37°C for about two hours or until single cell suspension was achieved. Every 15 minutes, the mixture was pipetted up and down with a cut P1000 pipette tip. Once single cell suspension was reached, 500 µl of inhibitor solution (supplied in the kit) was added to the well. The solution was gently mixed, filtered through a 70µm-pore sieve, and transferred to a 15 ml centrifuge tube. The cells were pelleted by centrifugation at 300 x g for 5 minutes at room temperature. Cell pellets were resuspended in 500µl of ice-cold 0.04% BSA diluted in PBS without Ca^2+^ and Mg^2+^. scRNA-seq was performed on 4-6 pooled 6-month-old organoids per donor line, per condition (hCS or hSS) for a total of 48 organoids. A minimum of 10,000 dissociated cells were submitted for sequencing. The library was prepared using the Chromium platform (10x Genomics) with the 3’ gene expression (3’ GEX) V3/V3.1 kit. Libraries were sequenced on an Illumina NovaSeq sequencer with an S4 flow cell, targeting a minimum depth of 20,000 reads per cell. The lengths for read parameters Read 1, i7 index, i5 index, and Read 2 were 100, 8, 0, 100, respectively.

#### vi. Bioinformatic analysis of scRNASeq data

The raw sequencing data, represented as base call (BCL) files produced by the Illumina sequencing platform, were subjected to demultiplexing and subsequent conversion into FASTQ format using CellRanger software (version 6.0.0, 10x Genomics) with default parameters. The software then mapped the FASTQ reads to the human reference genome (GRCh38) with default parameters. Following this, the ‘count’ command in the Cell Ranger (v6.0.0) software was utilized for the quantification of gene expression. For alignment and counting, the reference genome refdata-gex-GRCh38-2020-A was used, which was procured from the official 10x Genomics website. We performed QC and normalization using the Seurat (v3) R package^75^. For QC, we filtered out low-quality cells using the following criteria: (i) cells with unique gene counts outside the range of 200 to 6000; (ii) cells with more than 30% mitochondrial gene content; and cells with less than 2000 unique molecular identifiers^76,77^. Post QC, we carried out normalization, scaling the gene expression measurements for each cell by the total expression, multiplied by a scale factor (10,000 by default), and log-transformed the results. We extracted the expression profiles of the 338 genes identified by Birey et al^33^, to reduce the dimensionality of the dataset through principal component analysis (PCA), and identify statistically significant PCs using a JackStraw permutation test. This was followed by non-linear dimensional reduction using the UMAP (Uniform Manifold Approximation and Projection) technique for visualization. Cells were clustered based on their PCA scores using the Shared Nearest Neighbor (SNN) modularity optimization-based clustering algorithm in Seurat. After dimensionality reduction, we used known marker genes to guide the clustering of cells. Each cluster was then annotated using cell type markers identified by Birey et al^33^. To account for variability among individual donors we pooled genotypes, similar to our strategies in all other experiments, and performed a quasibinomial regression model to test if cell frequencies significantly differed between genotypes. Finally, we conducted differential expression analysis across the defined cell clusters using the FindAllMarkers function in Seurat, which employs a Wilcoxon Rank-Sum test to control for the variability within and between groups. The significantly differentially expressed genes were then used to interpret the biological significance of cell clusters. For each identified cell type, we conducted an enrichment analysis using the WebGestalt (WEB-based GEne SeT AnaLysis Toolkit) online tool, with the Human genome (GRCh38) as the reference set, employing a hypergeometric statistical method and the Benjamini & Hochberg method for multiple test adjustment.

#### vii. Analysis of ChIP-sequencing data from β-estradiol treated rodent brains

Tracks in bigWig format were downloaded from the GEO dataset GSE144718^40^. Two peaks within the *NRXN1* gene region in the mm10 genome were visualized using Spark (v2.6.2)^78^

#### viii. Generation of long-read sequencing data from post-mortem brain tissue

All aspects of sample processing (tissue handing, RNA extraction, library prep, QC, sequencing and bioinformatic analysis) was performed as previously described^15^.

### Therapeutic Treatments of iNeurons

#### i. β-estradiol treatment

β-estradiol was reconstituted in DMSO and subsequently diluted in Brainphys maturation media for a final concentration of 10 or 30nM. Neurons were treated for 3-5 consecutive days, with fresh drug or vehicle control replenished daily (in half media changes). iGLUT and iGABA neurons were treated starting from ∼DIV14-18. On the final day of treatment, cells were harvested ∼4 hours post dosage. Chronic treatments for functional experiments extended daily β-estradiol replenishments for ∼14 days and withdrawn 2-3 days prior to patch-clamp recordings. For RNASeq, the same chronic paradigm was performed and harvested on DIV21 ∼4 hours post final dosage.

#### ii. Antisense oligonucleotide treatment

A single HPLC-grade ASO was designed from Qiagen (LNA GapmeR) against the mutant (Ex 20/24) splice junction containing a phosphorothioate modified backbone with or without a 5’-FAM label for fluorescent visualization. All experiments were performed matched with a non-targeting ASO as the control group. ASOs were delivered using Lipofectamine RNAiMAX Transfection Reagent (Thermo, #13778075). The Lipofectamine RNAiMAX was diluted in Opti-MEM Medium (Thermo, #31985070). ASO was then diluted in Opti-MEM Medium at 1µM. The diluted ASO was added in a 1:1 ratio to the diluted Lipofectamine RNAiMAX and incubated at room temperature for 5 minutes. The ASO-lipid complex was added to cells, and incubated for 72 hours post-transfection, until RNA harvest. ASO samples for RNA-Seq were harvested on DIV17.

### Molecular Biology and Imaging

#### i. RNA extraction and RT-qPCR

For the isolation of RNA, 2D cells were lysed using TRIzol (Life Technologies #15596026) and purified using the Qiagen miRNeasy Kit (Qiagen Cat# 74106) according to the manufacturer’s instructions. For 3D organoids, pooled (early timepoints) or single organoids were washed and lysed using TRIzol, by manual homogenization with a pestle in a 1.5mL centrifuge tube. Following purification, samples were normalized within each experiment (15-50ng) and subsequently used for RT-qPCR assays using the *Power* SYBR Green RNA-to-Ct 1-Step Kit (Thermo REF 4389986). Relative transcript abundance was determined using the ΔΔCT method and normalized to the *ACTB* housekeeping gene. All primer sequences are listed below.

shRNA, primers and oligonucleotide probe sequences

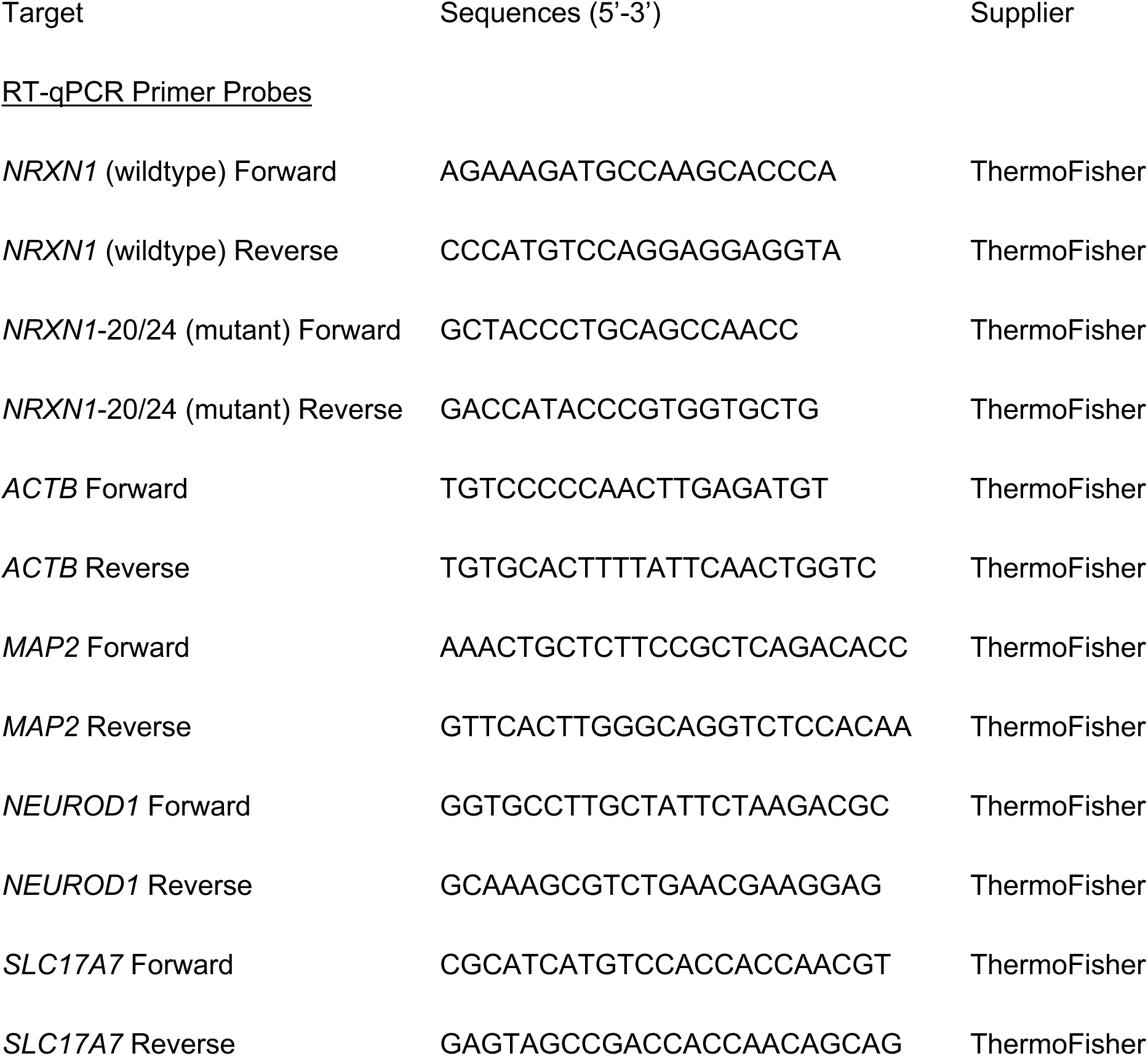
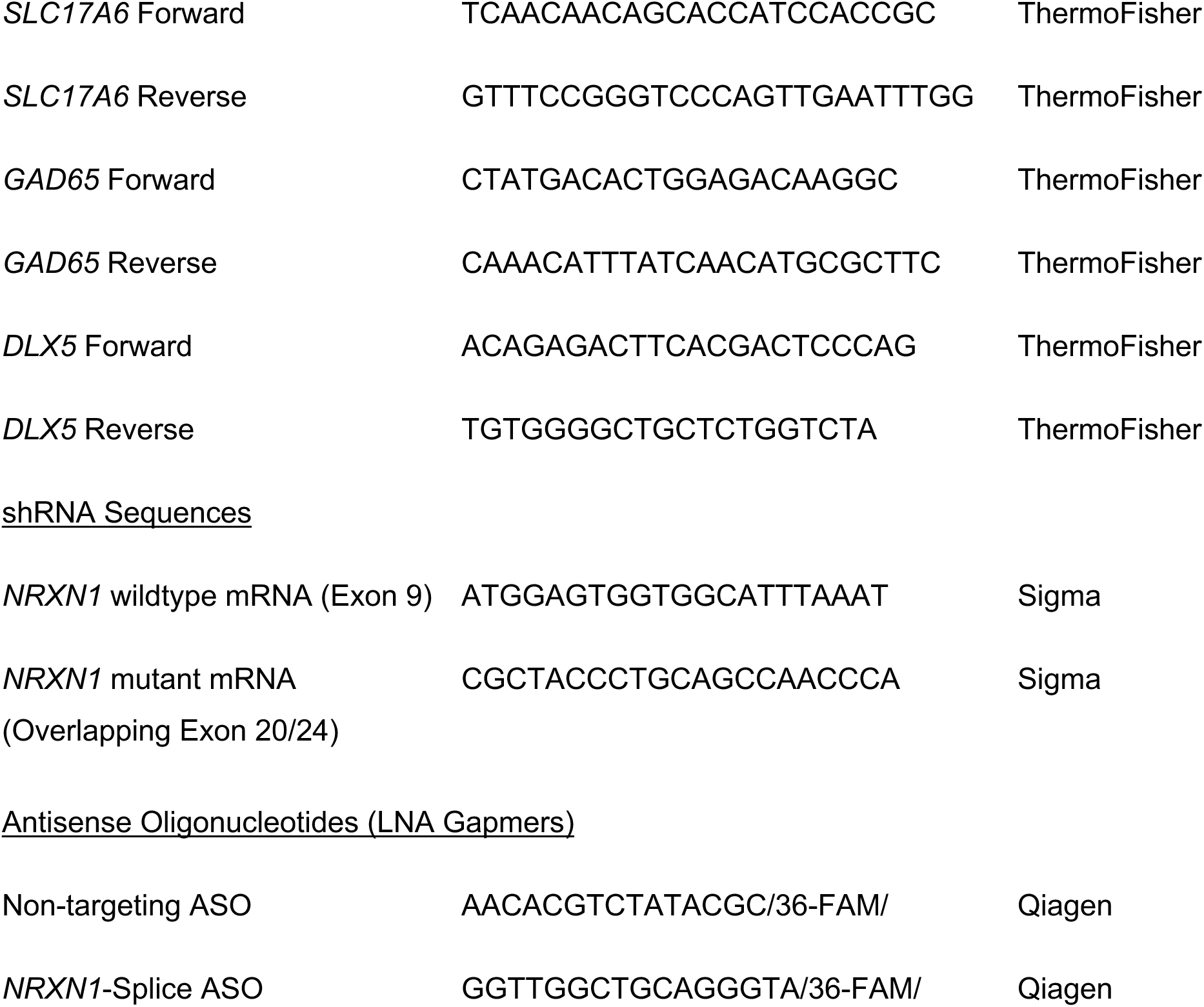

#### ii. Bright Field Imaging

For organoid perimeter analyses, brightfield microscope images of organoids were taken with a 2x objective. Image analysis was performed in ImageJ, with best fitting ovals or ellipses were selected around an organoid, and perimeter was measured.

#### iii. Immunostaining and imaging 2D cultures

For immunostaining of 2D monocultures, iGLUT and iGABA neurons seeded on acid-etched coverslips coated with PEI buffered with boric acid and 4x Matrigel. Samples were washed with DPBS Ca^2+/^Mg2^+^ and fixed using cold, fresh 16% paraformaldehyde (PFA, (Life Technologies, #28908)), diluted to 4% for 12 minutes at RT. Coverslips were then blocked and permeabilized with 2% donkey serum in DPBS Ca^2+/^Mg2^+^ supplemented with 0.1% Triton-X (Sigma, #93443-100ML) (blocking buffer), for one hour at RT. Primary antibody solutions were prepared in blocking buffer and incubated overnight at 4°C. The following day, samples were washed three times with PBS, and treated with secondary antibodies diluted in blocking buffer, for 1 hour in a dark chamber. Finally, samples were washed three times, and stained with DAPI for 15min at RT during the final wash. Coverslips were mounted with antifade (Vectashield #H-1000-10) onto glass slides and stored at 4°C until imaging using an upright Zeiss LSM 780 confocal microscope.

#### iv. High content imaging of 2D cultures

For high-content imaging, 100k cells/well were plated in an optically clear olefin bottom 96-well plate (Revvity Health Sciences, #6055302). At WPI4 (iGLUT) and WPI5 (iGABA), cultures were double fixed in PFA and 100% ice-cold methanol for 10min each with at least 2 washes of DPBS Ca^2+/^Mg2^+^ between and after fixation steps. Fixed cultures were washed twice in PBS and permeabilized and blocked two hours, followed by incubation with primary antibody solution overnight at 4°C. Cultures were then washed 3 times with PBS and incubated with secondary antibody solution for 1 hour at RT. Cultures were washed a further 3 timess with PBS with the second wash containing 1μg/ml DAPI. Fixed cultures were then imaged on a CellInsight CX7 HCS Platform with a 20x objective (0.4 NA) and neurite tracing and synaptic puncta detection performed using the synaptogenesis module in the Thermo Scientific HCS Studio 4.0 Cell Analysis Software to determine Syn1+ puncta density per µm of Map2+ve neurite length. 10 wells were imaged per donor with 9 images acquired per well for neurite tracing analysis. Wells with <10 annotated synapses, were excluded from the analysis.

| Target | Antibody Dilution | Supplier/ CAT# |
| --- | --- | --- |
| <u>Primary Antibodies</u> |  |  |
| (Rabbit) vGAT | 1:500 | Synaptic Systems, #135-303 |
| (Rabbit) vGLUT1 | 1:500 | Synaptic Systems, #131-002 |
| (Chicken) MAP2 | 1:1000 | Abcam, #ab5392 |
| (Mouse) Synapsin-1 | 1:500 | Synaptic Systems, #106 011 |
| DAPI | 1:1000 | Sigma, #D9542 |

| <u>Secondary Antibodies</u> |  |  |
| --- | --- | --- |
| (Donkey) 488-Mouse | 1:200 | Jackson ImmunoResearch, # 711-545-152 |
| (Donkey) 568-Rabbit | 1:200 | Abcam, #ab175700 |
| (Donkey) 647-Chicken | 1:500 | Life technologies, #A10042 |
| (Donkey) 568-Mouse | 1:500 | Life technologies, #A10037 |
| (Donkey) 488-Rabbit | 1:1000 | Life Technologies, #A-21206 |

### Statistics

No statistical power estimation analyses were used to predetermine sample sizes, where were chosen to match previous publications^14,21^ and field standards. All experimental statistics were performed in Prism *v10.1.1* and R *v4.2.0.* Bioinformatic analyses were performed in R *v3.5.3* (Bulk RNASeq DEG), *v4.1.2* (Bulk RNASeq GO) and *v4.1.2* (scRNAseq).

## Data and Code Availability

All source donor hiPSCs have been deposited at the Rutgers University Cell and DNA Repository (study 160; http://www.nimhstemcells.org/) and all bulk and single-cell transcriptome sequencing data are being prepared for deposits to GEO. To facilitate improved reproducibility of our data analyses, custom scripts has been deposited to github (https://github.com/mbfernando/NRXN1). Source data will be provided with this manuscript.

## Acknowledgements

MBF was supported by a Gilliam Fellowship from the Howard Hughes Medical Institute. This work was supported by the National Institute of Mental Health grants RO1 MH121074-02 (KJB, GF and PAS) and RO1 MH125579-02 (GF and KJB). DAK was supported by the National Science Foundation under Grant No. DBI2146398. Any opinions, findings, and conclusions or recommendations expressed in this material are those of the authors and do not necessarily reflect the views of the National Science Foundation. We thank the Stem Cell Engineering Core at the Icahn School of Medicine at Mount Sinai. We are grateful to the labs of Nan Yang (Ruiqi Hu and Xiaoting Zhou) and Samuele Marro (Madel Durens) for assistance in primary glial preparations. We are especially thankful to Kayla G. Townsley and Mark G. Baxter for advice on statistical testing, and Liang Yang for critical advice on scRNASeq analyses. We acknowledge Daniel Weinberger and the Lieber Institute for Brain Development at Johns Hopkins School of Medicine, for sharing post-mortem materials. Finally, the authors thank all members of the Brennand, Slesinger and Fang labs for critical feedback and discussions throughout the course of this work.

## Author Information

MBF, SK, ANM, RO, CP, AP, SG performed and analyzed experiments supervised by PAS and KJB. YZ and YF performed bioinformatic analyses supervised by GF. AT performed bioinformatic analysis of alternative splicing estimates, supervised by DAK. SW produced virus for the generation of iGABA neurons, and MD performed high-content imaging experiments. EKF processed post-mortem tissue and generated long-read data. IAP assisted in statistical analyses. MBF, GF, PAS and KJB wrote the paper with input from all authors.

## Ethics declarations / Competing interest statement

All authors have no competing interests to declare.

## Extended Data Figures

**Extended Data Figure 1:**
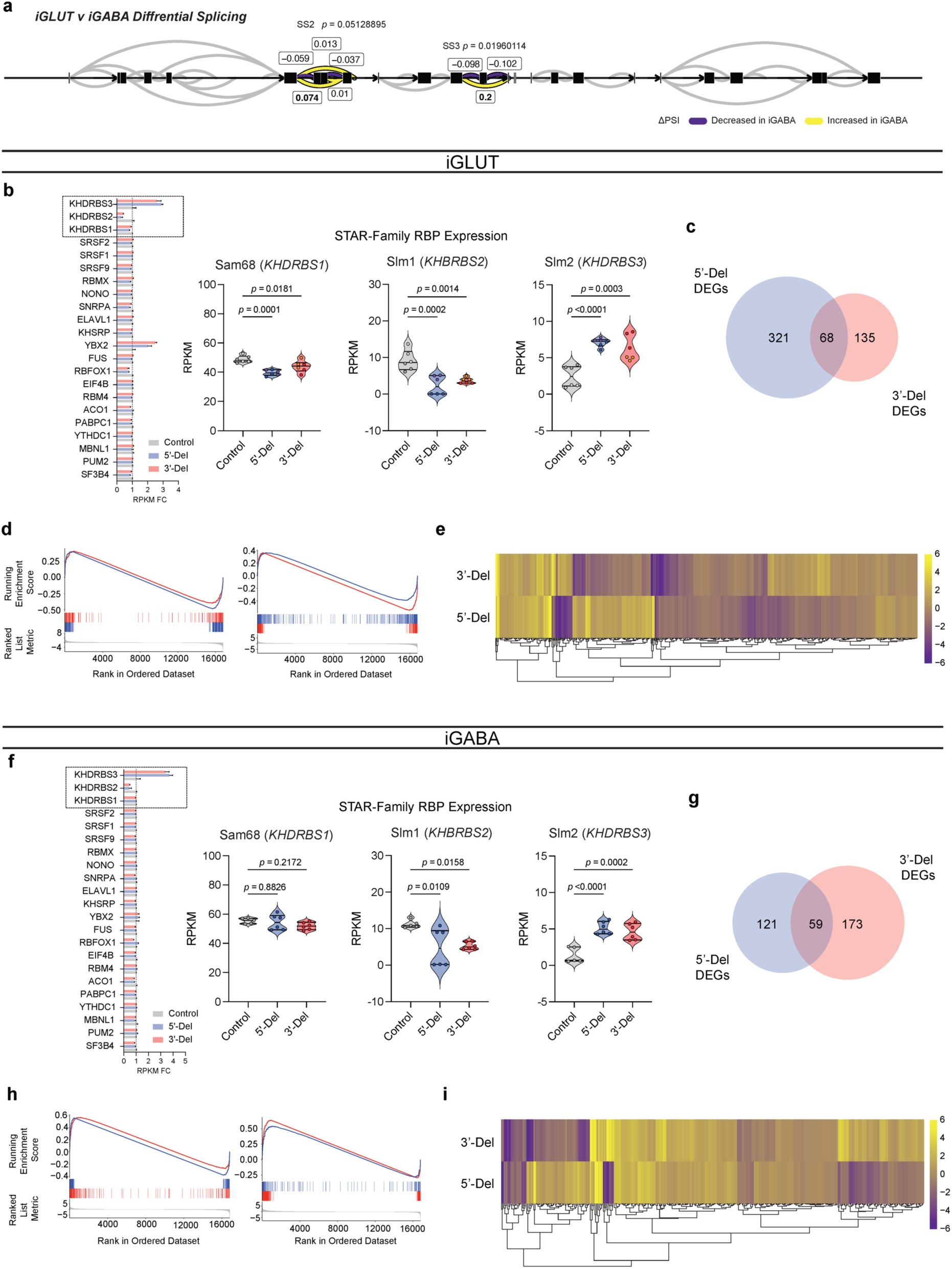
(**a**) Splicegraph displaying significant gene wide splicing clusters at *NRXN1* SS3 (*p* = 0.0196), compared via Dirichlet-multinomial generalized linear model with Bonferroni corrections. (**b, f**) Gene expression fold-change of select *NRXN1* predicted RNA-binding proteins (RBP) across patients and control iGLUT and iGABA neurons, with statistical comparisons for STAR-Family RBPs. (**c, g**) Overlap of DEGs and (**d,h)** gene set enrichment analysis (GSEA) between genotypes. (**e,i**) Distinct gene expression patterns by hierarchal clustering of all patient specific DEGs. Sample information correspond to Fig. 1

**Extended Data Figure 2:**
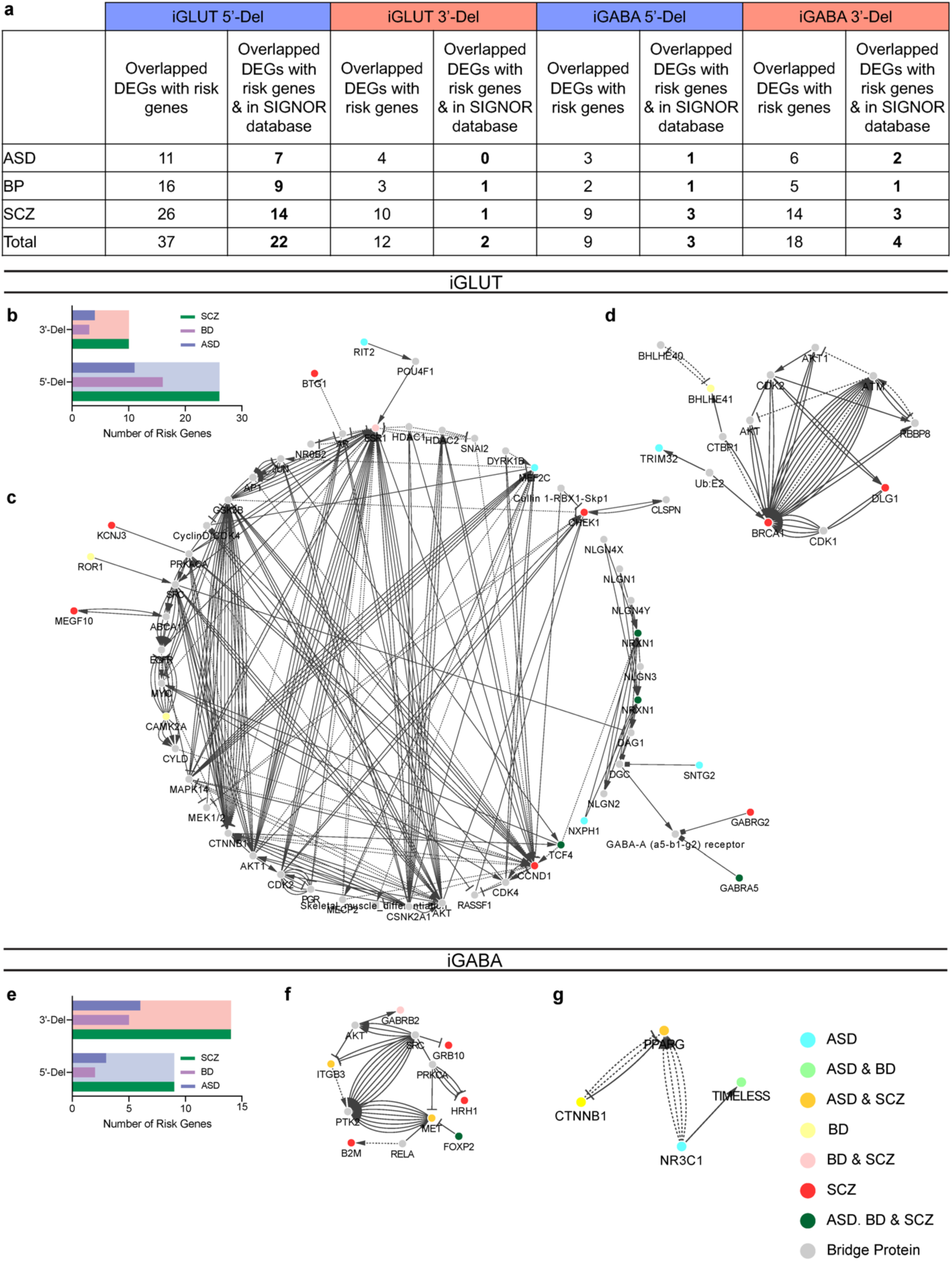
*Extended transcriptomics analysis on disease risk associated genes.* (a) Summary table of overlapping DEGs with risk enrichments across publicly curated datasets for autism (ASD), bipolar disorder (BD) and schizophrenia. (**b, e**) Enrichment of genes across neuropsychiatric disorders for iGLUT and iGABA neurons. (**c**) Interaction maps of risk genes for 5’-Del iGLUT, (**d**) 3’-Del iGLUT, (**f**) 5’-Del iGABA and (**g**) 5’-iGABA. Sample information correspond to Fig. 1

**Extended Data Figure 3:**
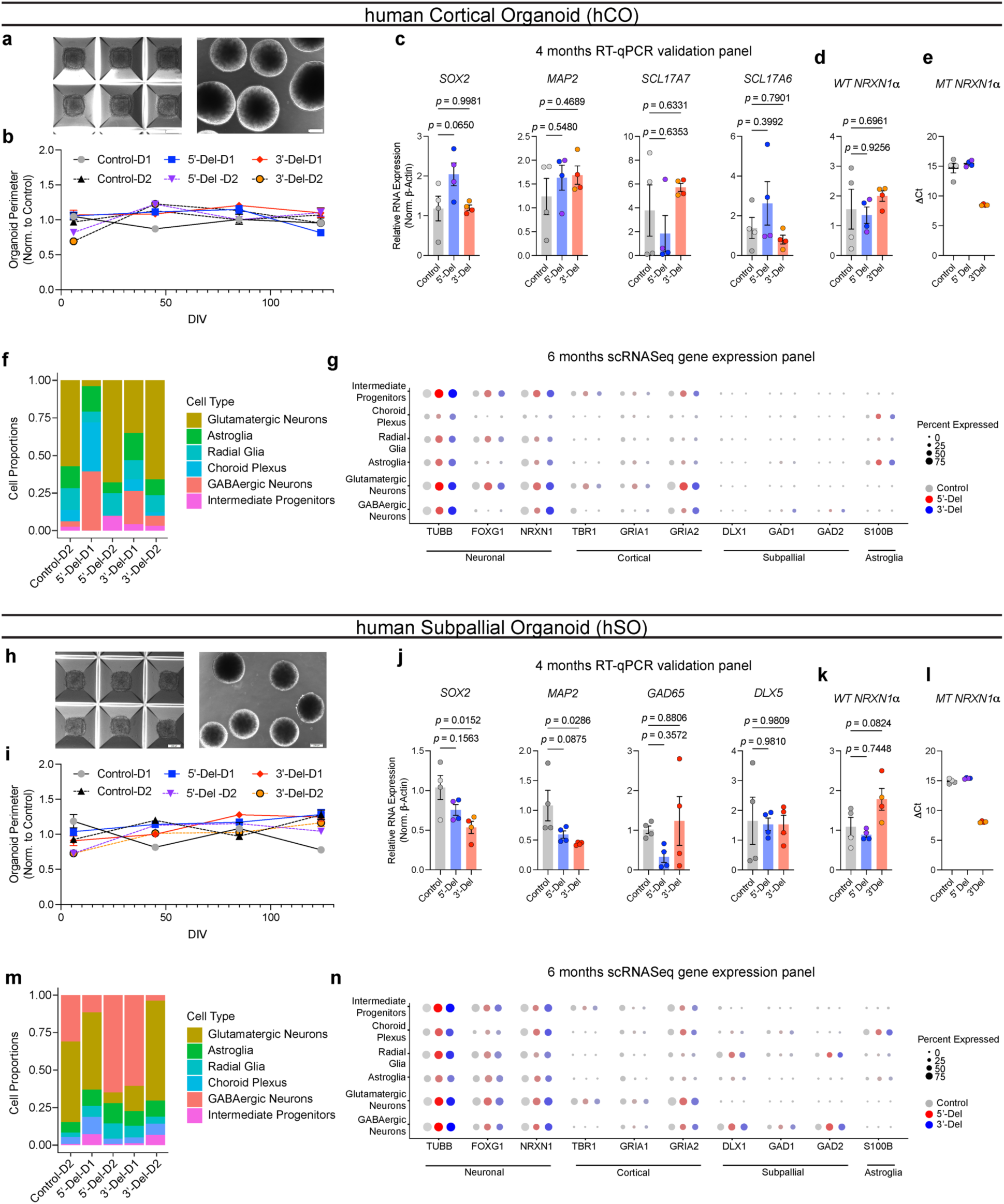
*Extended data on human organoid generation and characterization.* (**a,h**) Representative images of hiPSC aggregation and immature spheroids post dislodging. (**b,i**) Normalized organoid perimeters over time (compared to averaged control), hCO (n = 6 donors | 2 batches | 72-161 organoids) and hSO (n = 6 donors | 2 batches | 46-134 organoids). (**c, j**) RT-qPCR results from 4-month organoids of genes for pluripotency, neuronal, and cell-type specific markers. (d-e, k-l) RT-qPCR results of *NRXN1* WT and MT expression hCO (n = 6 donors | 1 representative batch | 12 samples) and hSO (n = 6 donors | 1 representative batch | 12 samples). Statistical tests used were 1-way ANOVAs with Dunnett’s test. (f, m) relative proportions of cell clusters across individual donors, (hCO = 47,460 cells) and (hSO = 35,563 cells). (**g, n**) Comprehensive gene expression panel across sub-clusters of hCO and hSO samples across neuronal, cortical, subpallial and astroglia markers. Data corresponds to Fig 2.

**Extended Data Figure 4:**
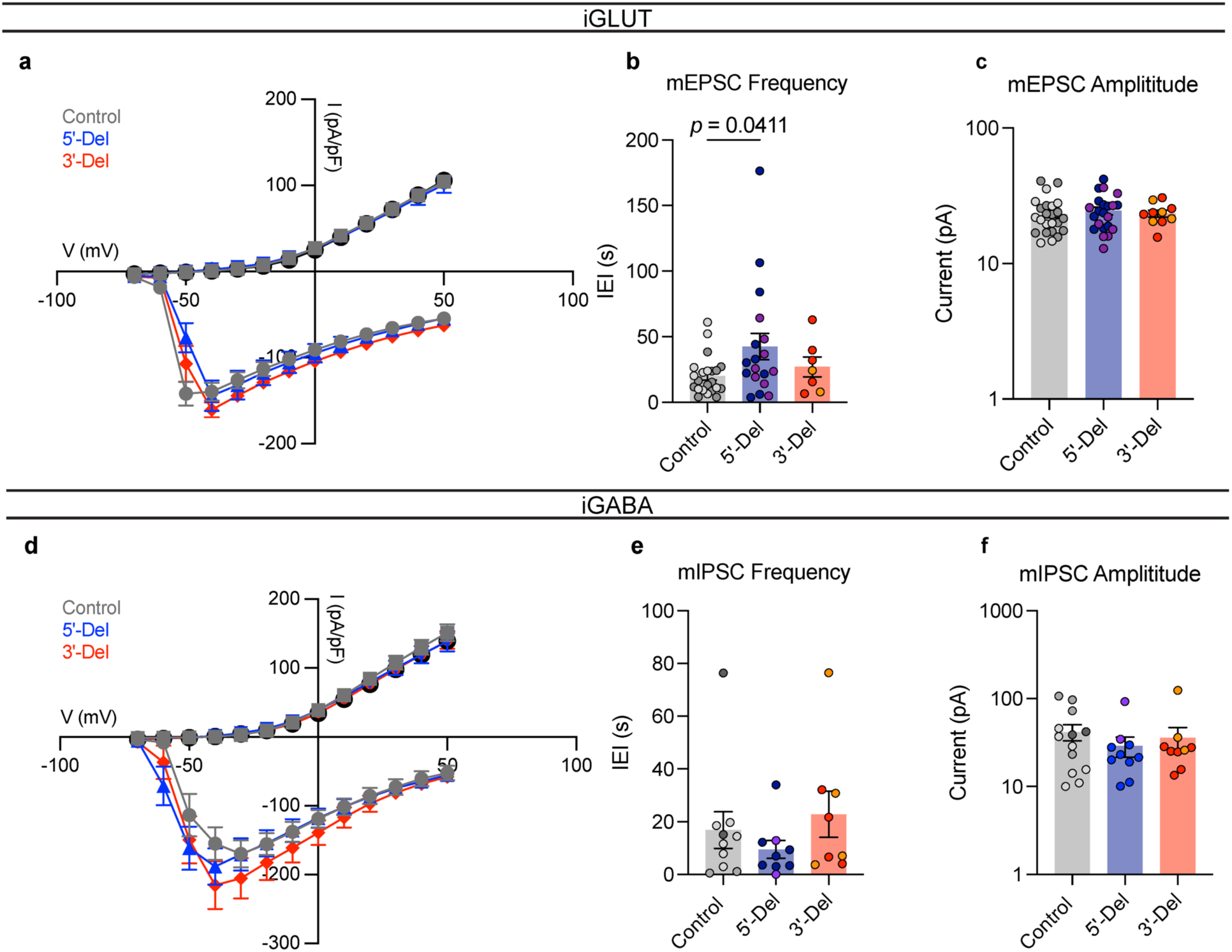
*Extended data on electrophysiological properties of 5’-Del and 3’-Del neurons.* (**a**) Voltage-gated potassium and channel kinetics across genotypes for iGLUT neurons (n = 6 donors | 2 inductions | 45 neurons). (**b**) Comparative mEPSC kinetics of IEI and (**c**) amplitude size from iGLUT neurons (n= 6 donors | 4 inductions | 47 neurons), compared via a 1-way ANOVA with Dunnett’s test. (**d**) Voltage-gated potassium and channel kinetics across genotypes for iGABA neurons (n = 6 donors | 2 inductions | 34 neurons). (**e**) Comparative mIPSC kinetics of (a) IEI and (**f**) amplitude size from iGABA neurons (n= 6 donors | 3 inductions | 27 neurons), compared via a 1-way ANOVA with Dunnett’s test. See Supplementary Table 5 for all summary statistics.

**Extended Data Figure 5:**
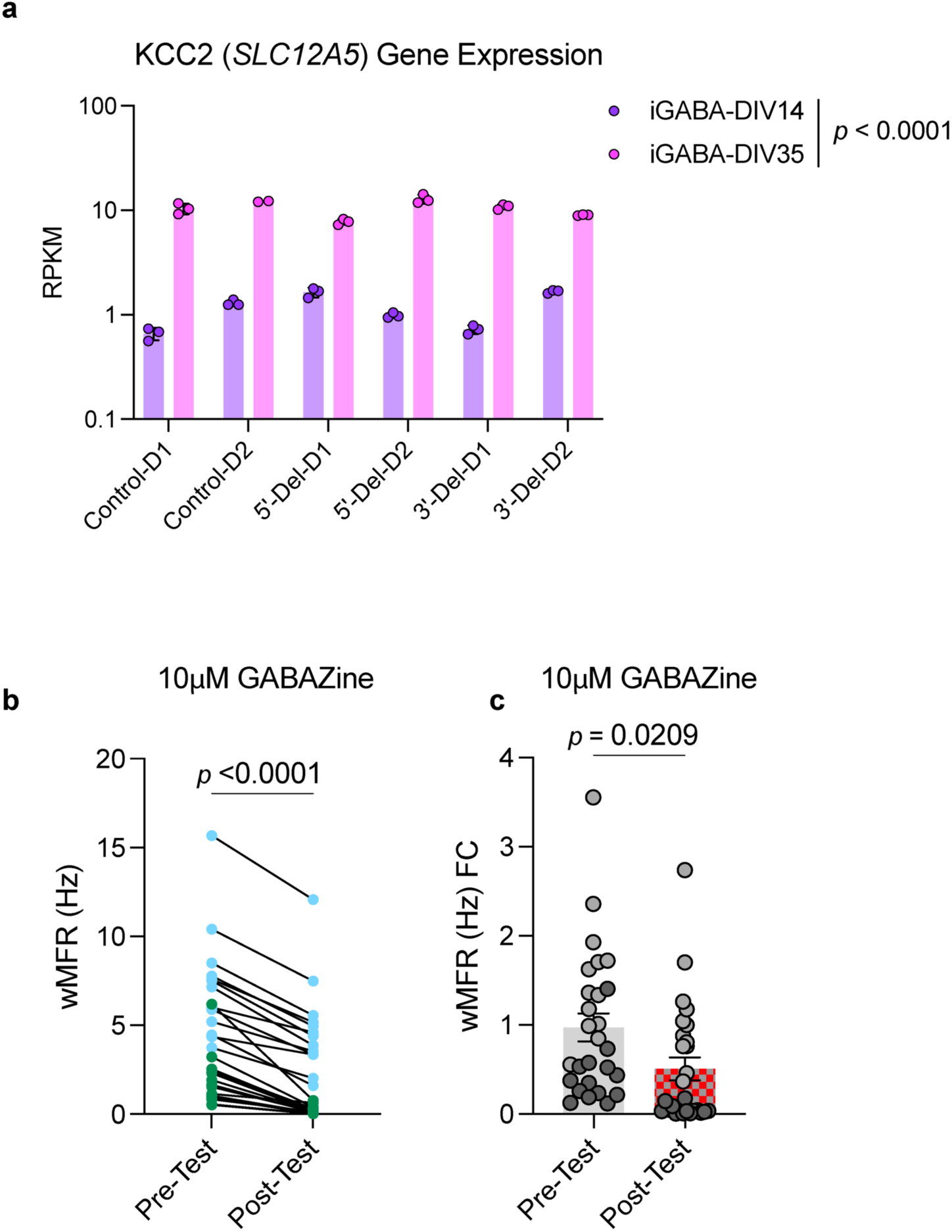
*Extended KCC2 related data from immature GABA neurons.* (a) Transcriptomic comparison of *SLC12A5* expression across DIV14 and DIV35 RNASeq timepoints. (b,c) MEA tests from pre- and post-treatment of 10uM GABAzine. (n = 2 donors | 1 representative induction | 28 MEA wells) Statistical tests are paired student’s t-test for time-linked comparison and unpaired student’s t-test for pre/post activity foldchange.

**Extended Data Figure 6:**
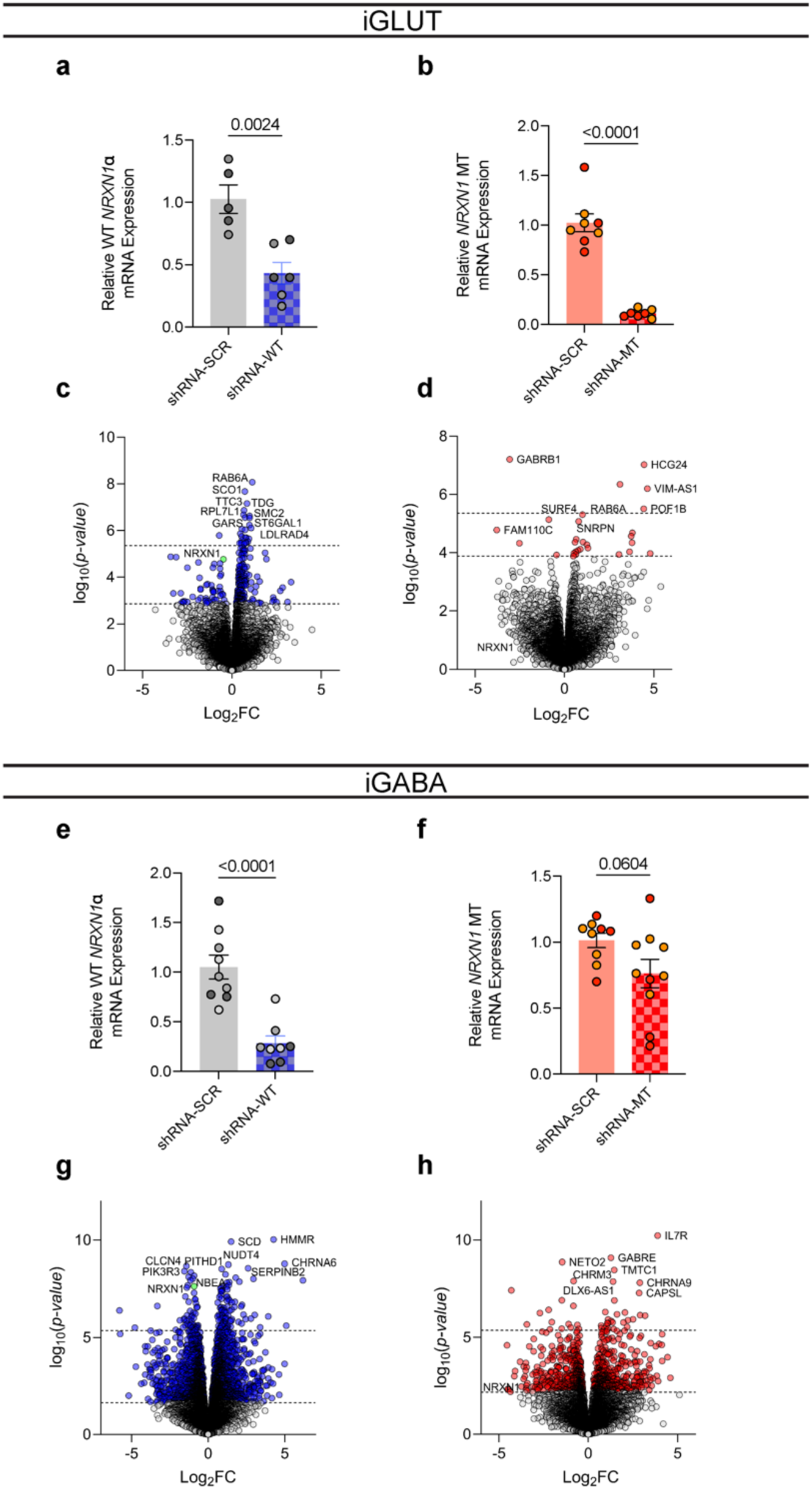
*shRNA knockdown validation.* (a) Extent of shRNA knockdown on WT and (b) MT *NRXN1* expression in iGLUT neurons (n = 2 donors | 1-3 inductions). (c) Extent of shRNA knockdown on WT and (d) MT *NRXN1* expression in iGABA neurons, (n = 2-3 donors | 1-3 inductions) Statistical tests used were Student’s t-test.

**Extended Data Figure 7:**
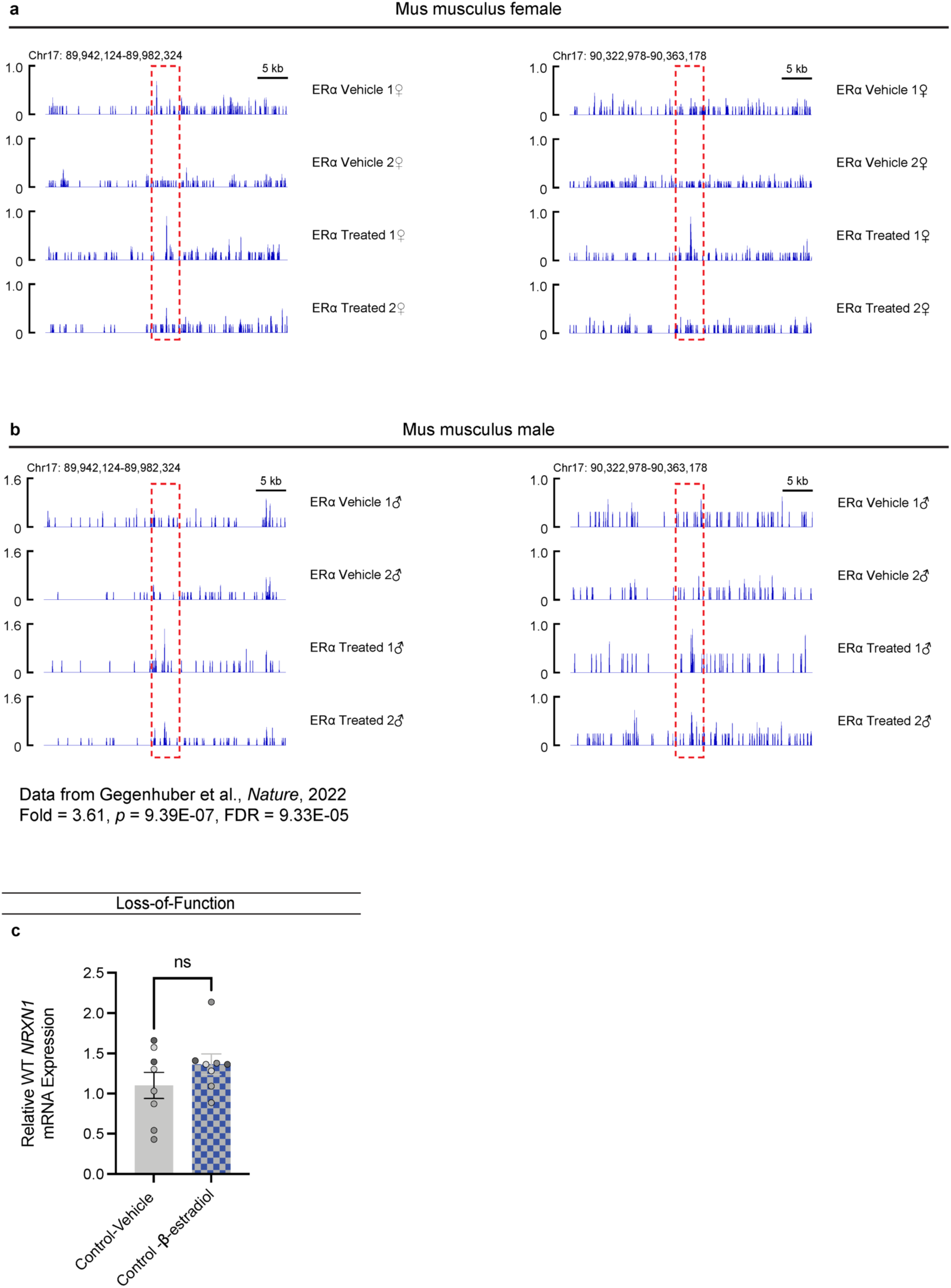
*ChIP-seq enrichment of ER1 binding at NRXN1 loci in rodent brain.* (**a**) Female and (**b**) male mus musculus ChIP tracts of *NRXN1* locus, with red dashed areas highlighting binding enrichment across vehicle and estradiol treated mice. (**c**) Effect of beta-estradiol on control donors (n = 16/4 | Representative).

**Extended Data Figure 8:**
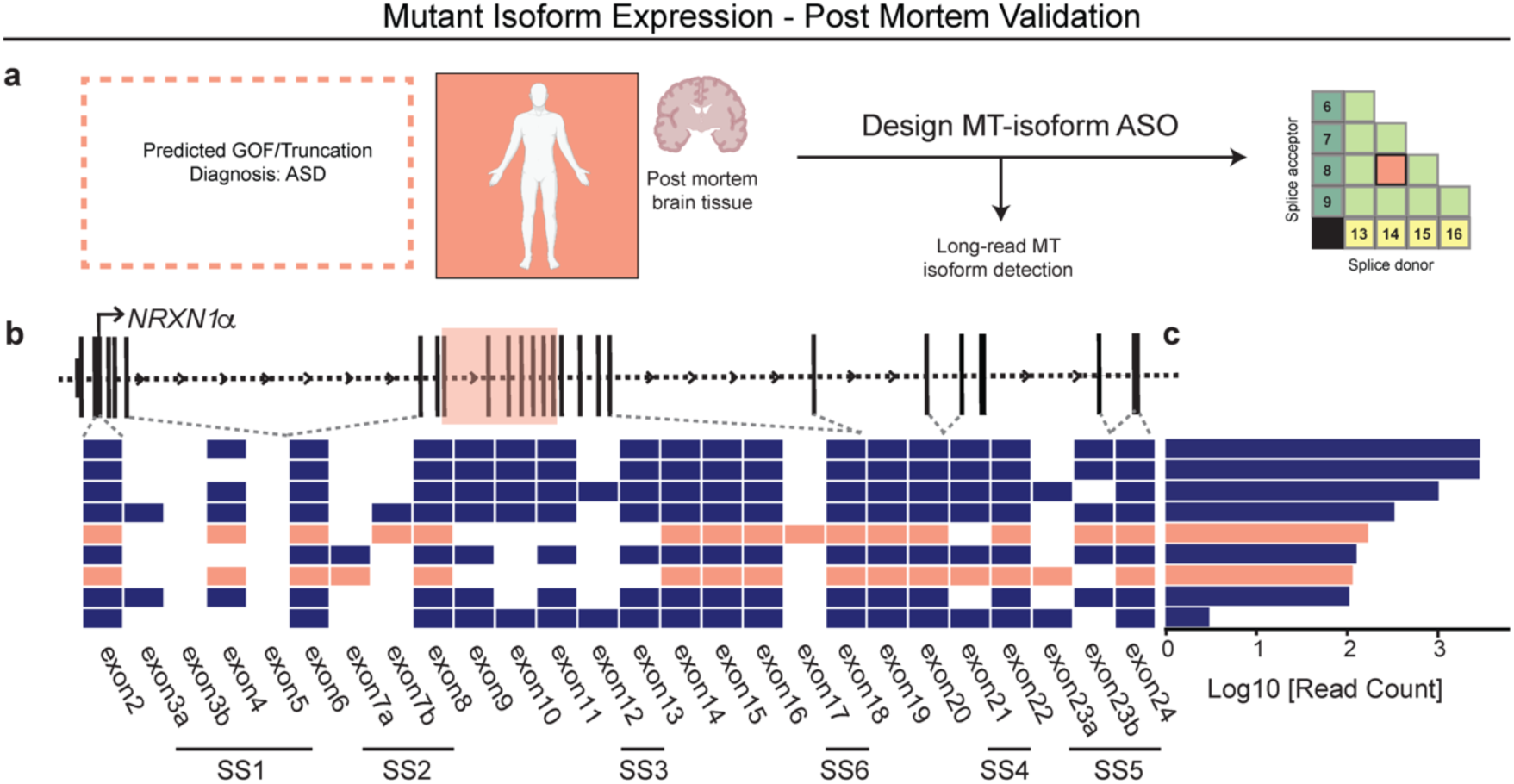
*In-vivo validation of MT isoform expression from an unrelated autism NRXN1^+/-^ patient.* (a) Schematic of novel NRXN1 autism patient, and GOF therapeutic targeting pipeline, with (b) schematic of the *NRXN1α* isoform structures, with each row representing a unique *NRXN1α* isoform and each column representing a *NRXN1* exon. The colored isoforms (navy, wildtype; peach, patient-specific) are spliced into the transcript while the blank exons are spliced out. (c) The abundance of each *NRXN1α* isoform by sample.

